# An energy-dissipative state of the major antenna complex of plants

**DOI:** 10.1101/2021.07.09.450705

**Authors:** Pierrick Bru, Collin J. Steen, Soomin Park, Cynthia L. Amstutz, Emily J. Sylak-Glassman, Lam Lam, Agnes Fekete, Martin J. Mueller, Fiamma Longoni, Graham R. Fleming, Krishna K. Niyogi, Alizée Malnoë

## Abstract

Plants and algae are faced with a conundrum: harvesting sufficient light to drive their metabolic needs while dissipating light in excess to prevent photodamage, a process known as non-photochemical quenching. A slowly relaxing form of energy dissipation, termed qH, is critical for plants’ survival under abiotic stress. Here, we tested whether we could isolate photosynthetic subcomplexes (from plants in which qH was induced) that would remain in an energy-dissipative state. Interestingly chlorophyll fluorescence lifetimes were decreased by qH in isolated major trimeric antenna complexes, providing a natively quenched complex with physiological relevance to natural conditions. Next, we monitored the changes in thylakoid pigment, protein or lipid content of antenna with active or inactive qH, and no evident differences were detected. Finally, we investigated whether specific antenna subunits of the major antenna were required for qH but found it insensitive to trimer composition. Because qH can occur in the absence of specific xanthophylls, and no changes in pigments were detected, we propose that the energy-dissipative state reported here may stem from chlorophyll-chlorophyll excitonic interaction.

## Introduction

Photosynthetic organisms possess pigment-protein antenna complexes, which can switch from a light-harvesting state to an energy-dissipating state (Valkunas et al., 2012, Liguori et al., 2015). This switching capability regulates how much light is directed towards photochemistry and ultimately how much carbon dioxide is fixed by photosynthesis (Zhu et al., 2010). The fine-tuning of light energy usage is achieved at the molecular level by proteins which act at or around these pigment-protein complexes (Demmig-Adams et al., 2014). Understanding the regulatory mechanisms involved in the protection against excess light, or photoprotection, has important implications for engineering optimized light-use efficiency in plants, and thereby increasing crop productivity and/or tolerance to photooxidative stress (Ort et al., 2015, Kromdijk et al., 2016, Wang et al., 2020).

Non-photochemical quenching (NPQ) processes protect photosynthetic organisms by safely dissipating excess absorbed light energy as heat (Horton et al., 1996, Müller et al., 2001). Several NPQ mechanisms have been described and classified based on their recovery kinetics and/or molecular players involved (Malnoë, 2018, Pinnola & Bassi, 2018, Bassi & Dall’Osto, 2021). In plants, the rapidly reversible NPQ (relaxes within minutes), qE, relies on ΔpH, the protein PsbS, and the xanthophyll pigment zeaxanthin (Niyogi & Truong, 2013). The slowly reversible NPQ (relaxes within hours to days), or sustained energy dissipation, includes several mechanisms such as qZ (zeaxanthin-dependent, ΔpH-independent (Dall’Osto et al., 2005)), qH (see below), and qI (due to photosystem II (PSII) reaction center subunit D1 photoinactivation (Krause et al., 1990) which can be reversed by D1 repair (Nawrocki et al., 2021)). Energy re-distribution can also cause a decrease, i.e. quenching, of chlorophyll (Chl) fluorescence through qT (due to state transition, movement of antenna phosphorylated by the kinase STN7 (Quick & Stitt, 1989)).

We have recently uncovered, using chemical mutagenesis and genetic screens in *Arabidopsis thaliana*, several molecular players regulating qH (Brooks et al., 2013, Malnoë et al., 2018, Amstutz et al., 2020, Bru et al., 2020). qH requires the plastid lipocalin, LCNP (Malnoë et al., 2018), is negatively regulated by suppressor of quenching 1, SOQ1 (Brooks et al., 2013), and is inactivated by relaxation of qH 1, ROQH1 (Amstutz et al., 2020). Importantly, qH is independent of PsbS, ΔpH, xanthophyll pigments and phosphorylation by STN7 (Brooks et al., 2013, Malnoë et al., 2018). Strikingly, when qH is constitutively active in a *soq1 roqh1* mutant, plants are severely light-limited and display a stunted phenotype (Amstutz et al., 2020). If qH cannot occur (as in an *lcnp* mutant), a higher amount of lipid peroxidation is observed, and plants are severely light-damaged under stress conditions such as cold temperature and high light (Levesque-Tremblay et al., 2009, Malnoë et al., 2018). Our present working hypothesis is that, under stress conditions, LCNP binds or modifies a molecule in the vicinity of or within the antenna proteins, thereby triggering a conformational change that converts antenna proteins from a light-harvesting to a dissipative state.

In wild-type (WT) *Arabidopsis* plants, qH occurs in response to cold and high light (Malnoë et al., 2018), whereas the *soq1* mutant can display qH under non-stress conditions upon a 10-min high light treatment (Brooks et al., 2013). In plants, the peripheral antenna of PSII is composed of pigment-binding, light-harvesting complex (Lhcb) proteins, which are divided into minor subunits (Lhcb4, 5, 6 or CP29, 26, 24, respectively) present as monomers and major subunits (Lhcb1, 2, 3) also referred to as LHCII, forming hetero- and homo-trimeric complexes associated to PSII in a strongly, moderately or loosely bound manner (Ballottari et al., 2012, Crepin & Caffarri, 2018). The pigments associated with the major and minor antenna complexes include Chls *a* and *b*, and xanthophylls such as lutein, violaxanthin, zeaxanthin, and neoxanthin (Jahns & Holzwarth, 2012). The mutant *chlorina1* does not accumulate trimeric Lhcbs because it lacks Chl *b*, but it does accumulate some monomeric Lhcbs with Chl *a* only (Takabayashi et al., 2011). qH is no longer observed in the double mutant *soq1 chlorina 1* (Malnoë et al., 2018), indicating that qH may require the trimeric antenna and/or Chl *b*. Here, we investigated whether qH remained active upon isolation of photosynthetic subcomplexes with the aim to narrow down the location of the qH quenching site. We measured the chlorophyll fluorescence lifetimes of intact leaves, isolated thylakoids, and isolated complexes from WT and several *Arabidopsis* mutants relating to qH under non-stress and stress conditions with active or inactive qH. Isolation of partly quenched LHCII directly from thylakoid membranes with active qH showed that qH can occur in the major trimeric LHCII complexes. Through genome editing and genetic crosses, we further demonstrated that qH does not rely on a specific major Lhcb subunit, suggesting that qH is not due to specific amino acid variation among Lhcb1, 2 and 3 (such as phosphorylation in Lhcb1 and 2, or presence of cysteine in Lhcb2.3 or aromatic residues in Lhcb3) and/or that compensation from other major Lhcb proteins may occur. Prior to this work, only a few studies had reported a quenched conformation of isolated LHCII trimers and in contrast to the native isolation reported here, quenching was achieved in vitro, after full solubilization of LHCII (van Oort et al., 2007, Ilioaia et al., 2008). Successful isolation of natively quenched LHCII paves the way for revealing its molecular origin.

## Results

### Chlorophyll fluorescence lifetimes are decreased by qH in leaves and thylakoids

Previously, we demonstrated that qH is induced by a cold and high light treatment on whole plants of *Arabidopsis* in mutants, and importantly also in wild type. We found that the amount of NPQ measured by chlorophyll fluorescence imaging can reach a high level, approximately 12 in the *soq1* mutant, and this induction of NPQ is LCNP-dependent as it does not occur in the *soq1 lcnp* double mutant (Malnoë et al., 2018). We also observed constitutive qH from non-treated plants in the *soq1 roqh1* double mutant which displayed maximum fluorescence (Fm) values ~85% lower than wild type or *soq1 roqh1 lcnp*, indicating a high NPQ yield (Amstutz et al., 2020). To ascertain that qH under stress condition such as cold and high light, or in the double mutant *soq1 roqh1*, was due to a decrease in chlorophyll excited-state lifetime, we measured fluorescence lifetime on both intact leaves and thylakoids via time-correlated single photon counting. Here, we used the laser at a saturating light intensity to close PSII reaction centers so that differences in lifetime can be attributed to NPQ (Sylak-Glassman et al., 2016). Strikingly, non-treated *soq1 roqh1* indeed displayed a decreased amplitude-weighted average fluorescence lifetimes (τ_avg_) compared to controls (Fig.1, light grey bars), with a much shorter value in both leaves (~0.1 ns vs. ~1.5 ns) and thylakoids (~0.2 ns vs. ~1.1 ns). These data unequivocally provide evidence that LCNP-dependent NPQ, qH, promotes a chlorophyll de-excitation pathway, which remains active upon isolation of thylakoid membranes.

**Figure 1.**
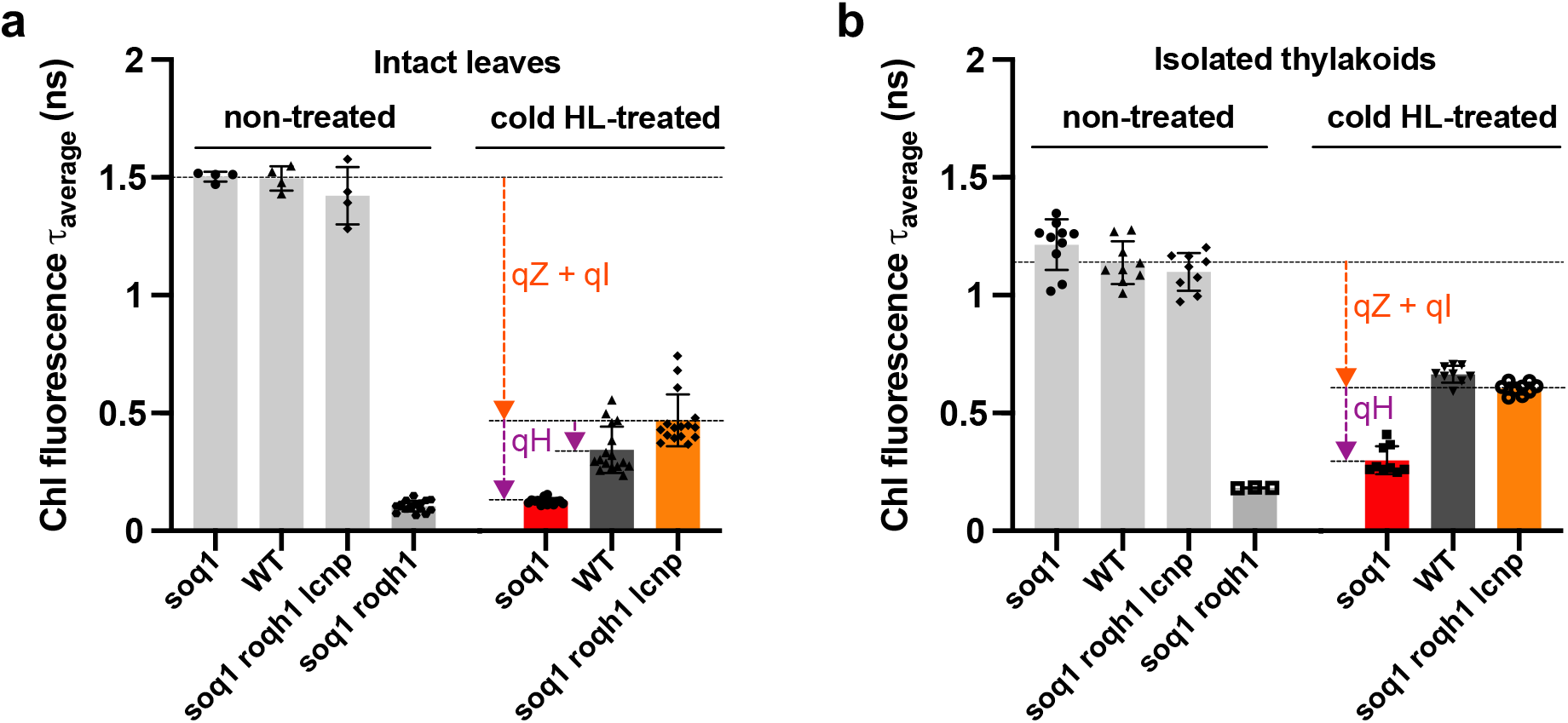
qH decreases chlorophyll fluorescence lifetimes of leaves and thylakoids. Average fluorescence lifetime (τ_average_) of intact leaves **(a)** or crude thylakoid membrane **(b)** from non-treated *soq1*, wild type (WT), *soq1 roqh1 lcnp* and *soq1 roqh1* or cold and high light (cold HL)-treated *soq1*, WT and *soq1 roqh1 lcnp* for 6 h at 6°C and 1500 μmol photons m^-2^ s^-1^. qE is relaxed by dark-acclimating for 5 min before each measurement (for non-treated isolated thylakoids, dark-acclimation of detached leaves overnight prior to thylakoid extraction). Excitation at 420 nm and emission at 680 nm. Data represent means ±SD (intact leaves, non-treated, n=4 plant individuals and n=17 for *soq1 roqh1*, cold HL-treated, n=14-16; isolated thylakoids, n=3-6 technical replicates from 2 independent biological experiments each with n > or = 5 plants; one biological experiment for *soq1 roqh1*). NPQτ values are determined based on: 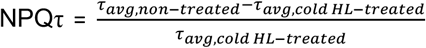. NPQτ of cold HL-treated leaves from *soq1*, WT and *soq1 roqh1 lcnp* are 11, 3.3 and 2, respectively and of cold HL-treated thylakoids 3.1, 0.7 and 0.8.

Next, we exposed plants to a 6-h cold and high light treatment (6°C and 1500 μmol photons m^-2^ s^-1^) followed by dark-acclimation for 5 min to relax qE. During this treatment, qH is induced and so is qZ as zeaxanthin accumulates (de-epoxidation state value of approximately 0.7 (stress) vs. 0.05 (non-stress) in all lines (Malnoë et al., 2018, Amstutz et al., 2020)); the remaining slowly relaxing quenching processes are grouped under the term qI and are in part due to photoinactivation of PSII. In Fig.1, colored bars show that the cold and high light-treatment results in significantly decreased chlorophyll excited-state lifetime in both leaves and isolated thylakoids. Interestingly, cold and high light-treated *soq1* leaves displayed τ_avg_ values similar to non-treated *soq1 roqh1* indicating that this stress treatment triggers a fully active qH state in *soq1*. Wild-type leaves displayed an intermediate τ_avg_ value between active qH (*soq1*) and inactive qH (*soq1 roqh1 lcnp*) further establishing the occurrence, and physiological relevance, of qH in promoting energy dissipation under abiotic stress (Fig. 1a). The calculated NPQτ derived from the τ_avg_ values were all in agreement with the NPQ value measured by pulse-amplitude modulated chlorophyll fluorescence in (Malnoë et al., 2018, Amstutz et al., 2020). We also measured the fluorescence lifetime from leaves of *soq1 npq4 roqh1* (lacks SOQ1, PsbS and ROQH1), *soq1 roqh1* ROQH1 overexpressor (OE) and several mutant alleles of *roqh1* and *soq1 roqh1* (Fig. S1, Table S1). The τ_avg_ values, and calculated NPQτ, further highlighted that qH is independent of PsbS (*soq1 npq4 roqh1* has a low τ_avg_ in a similar range to *soq1 roqh1*, ~60 ps vs. 130 ps) and that ROQH1 is required for relaxation of qH (NPQτ of *roqh1* mutant and *soq1 roqh1* ROQH1 OE, respectively higher and lower than wild type).

Of note, in cold and high light-treated thylakoids, the τ_avg_ values were overall higher than observed in intact leaves, and the difference in τ_avg_ between wild type and *soq1 roqh1 lcnp* was no longer apparent (Fig. 1b). We probed the release of chlorophyll fluorescence by step solubilization of thylakoid membrane preparation (Fig. S2) and it showed that qH is partly due to protein–protein and lipid–proteins interactions in the membrane (cold and high light, Q_m_ *soq1* higher than *soq1 roqh1 lcnp*) and due to pigment–protein interactions (Q_pi_ also higher) which may explain the lower NPQτ of *soq1* thylakoids compared to leaves as some of these interactions may have been lost during thylakoid preparation. Yet, although smaller, the retention of active qH in *soq1* thylakoids offered a unique opportunity to explore whether quenched photosynthetic subcomplexes could be isolated.

### Isolated LHCII trimers from plants with active qH are quenched

Next, we tested whether we could observe qH in a specific isolated pigment-protein complex. The lines *soq1* (active qH) and *soq1 lcnp* (inactive qH) were chosen for this purpose (*soq1 roqh1* and *soq1 roqh1 lcnp* could have been used but *soq1 roqh1* plants are much smaller due to light limitation by constitutive active qH). Plants were treated with cold and high light for 6 h at 6°C and 1500 μmol photons m^-2^ s^-1^ followed by dark-acclimation for 5 min to relax qE. Thylakoids were isolated, solubilized, and fractionated by gel filtration to separate complexes based on their size. The separation profiles of photosynthetic complexes were similar for *soq1* and *soq1 lcnp* (Fig. S3a). Fractions corresponding to PSII-LHCII mega-complexes, supercomplexes, PSI-LHCI supercomplexes, PSII core dimer, LHCII trimer, and LHCII/Lhcb monomer as well as smaller fractions (peaks 7, 8) were collected, and their fluorescence yield was measured by video imaging (Fig. S3b). The LHCII trimer fraction clearly displayed a relatively lower fluorescence yield with active qH. Room-temperature fluorescence spectra were measured at the same low Chl concentration (0.1 μg mL^-1^) to prevent re-absorption and with excitation at 625 nm (isosbestic point) to excite both Chls *a* and *b* equally; the Chl *a/b* ratio is similar between the compared samples so absorption at 625 nm should be equal. Complexes from non-treated wild type were isolated for reference; material came from plants grown under standard light conditions. The LHCII trimer fraction displayed a fluorescence yield at 680 nm that was on average 24% ± 8% lower with active qH compared to inactive qH and WT reference, whereas the LHCII/Lhcb monomer fraction displayed no significant differences among samples (Fig. 2a, Fig. S4a,b). A complementary approach separating pigment-protein complexes following solubilization by clear native-PAGE further evidenced that qH is active in isolated LHCII trimers (Fig. 2b, Fig. S4c). These results suggest that qH occurs in the LHCII trimer and remains active even after isolation of the solubilized protein complex.

**Figure 2.**
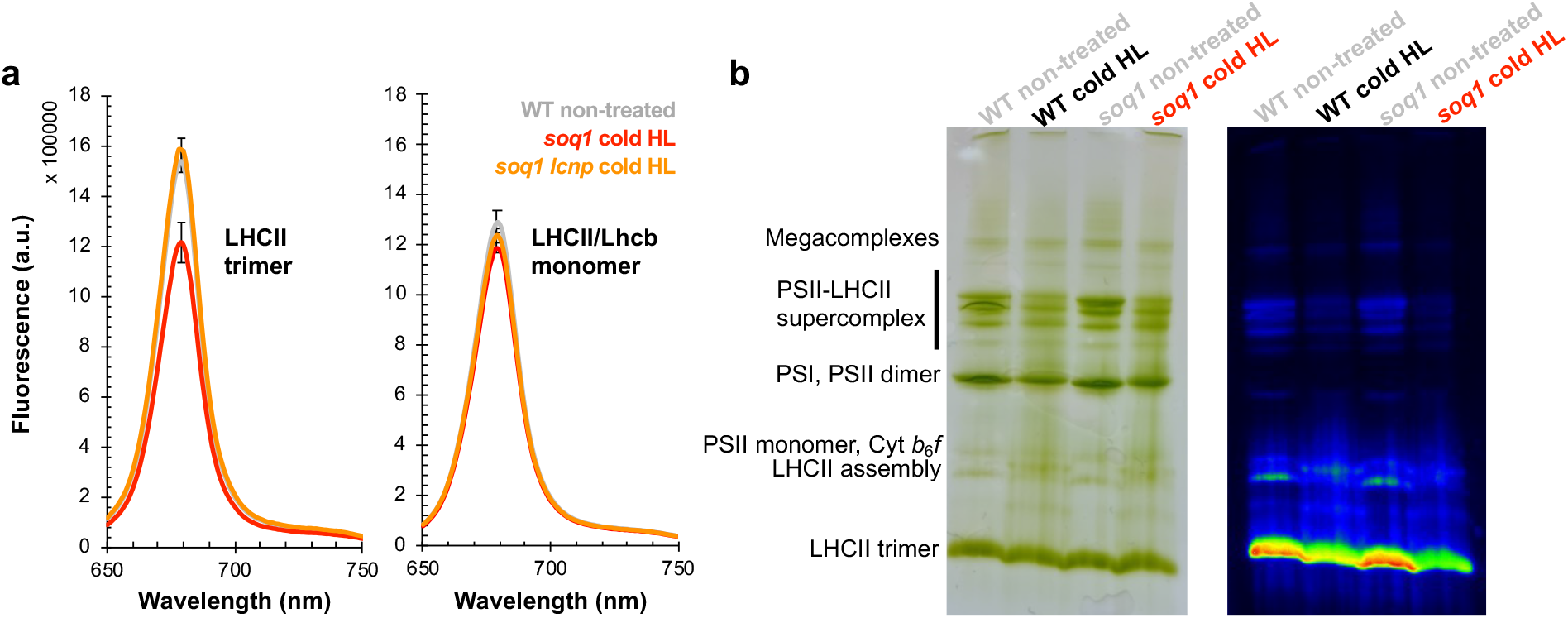
Isolated LHCII trimers from plants with active qH are quenched. **(a)** Room temperature fluorescence spectra of isolated LHCII trimer (left) and LHCII/Lhcb monomer (right) pooled fractions from non-treated wild type (WT) (grey) and cold and high light (cold HL)-treated *soq1* (red) and *soq1 lcnp* (orange) (see Fig. S3 for gel filtration experiments and peaks annotation from which fractions were pooled). Fluorescence emission from 650 nm to 750 nm from samples diluted at same chlorophyll concentration (0.1 μg mL^-1^) with excitation at 625 nm, with maxima at 679 nm for all samples. Data represent means ±SD (n=3 technical replicates from biological replicate 3 with n=8 plants). Representative from three independent biological experiments is shown (see Fig. S4a,b for biological replicates 1 and 2). **(b)** Thylakoids were extracted from WT and *soq1* plants (n=3 individuals of each) grown under standard conditions (grey) or cold HL-treated for 10 h (black and red) with NPQ values of respectively 3±1 and 11±1, solubilized in 1% α-DM and separated by clear native PAGE on a 3-12% gel. 10 μg Chl were loaded per lane. Gel image (left) and chlorophyll fluorescence image (right). The composition of the major bands are indicated based on (Rantala *et al.* (2018)). Representative from two independent biological experiments (with n > or = 3 plants) is shown (see Fig.S4c for LHCII trimer pigment-protein band and chlorophyll fluorescence quantification).

We measured the chlorophyll fluorescence lifetimes of LHCII trimer, LHCII/Lhcb monomer and PSII dimer fractions. We observed in the active qH LHCII trimer fraction a ~20% shorter τ_avg_ compared to that of inactive qH (~2.6 ns for *soq1* vs. ~3.3 ns for *soq1 lcnp*) in agreement with the ~20% decrease in yield (Fig. 3); for reference non-treated WT LHCII τ_avg_ is ~3.1 ns (Table S2). No differences in fluorescence lifetimes due to qH were detected in either the LHCII/Lhcb monomer or PSII dimer fractions. These results demonstrate that qH promotes a chlorophyll deexcitation pathway in the trimeric antenna, and is distinct from qI.

**Figure 3.**
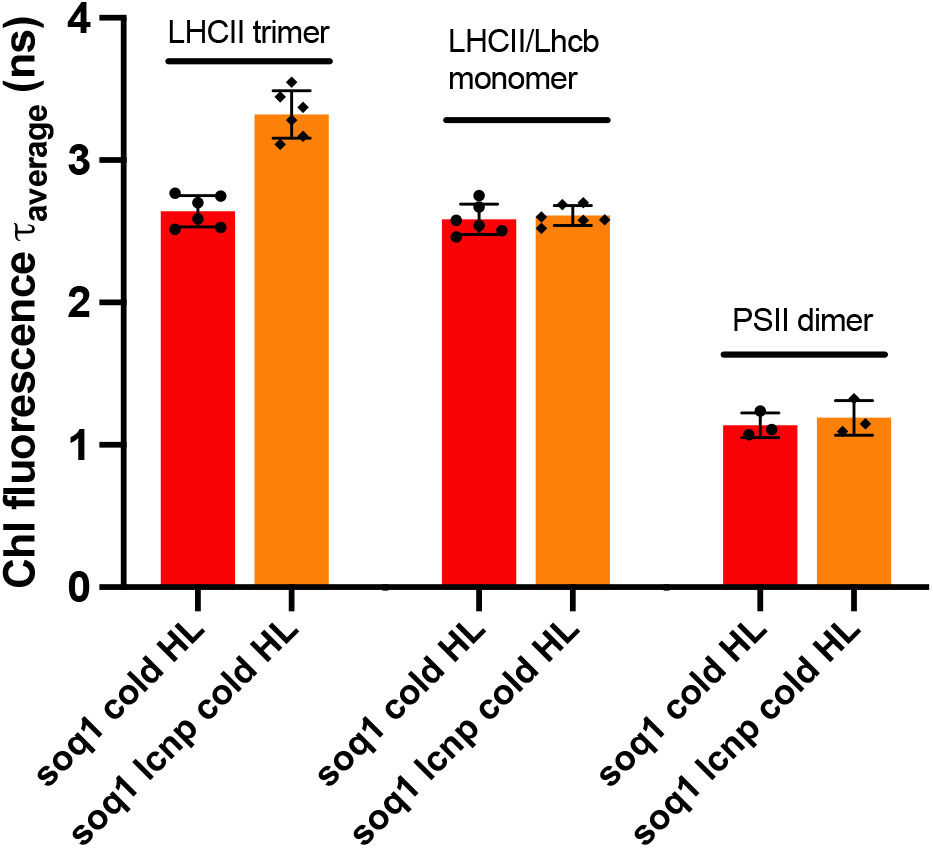
qH decreases chlorophyll fluorescence lifetimes of isolated LHCII trimers. Average fluorescence lifetime (τ_avg_) of LHCII trimer, LHCII/Lhcb monomer, and PSII dimer isolated from cold HL-treated *soq1* (red) and *soq1 lcnp* (orange). Data represent means ±SD (n=3 technical replicates from 2 independent biological experiments each with n = 8 plants).

### No evident changes in pigment, lipid and protein content of quenched LHCII

We examined the pigment, lipid and protein content by HPLC, LC-MS and SDS-PAGE, respectively, to investigate which molecular changes may be at the origin of the qH-energy dissipative state in the trimeric antenna. There were no apparent differences in pigment composition (Fig. S5a) or abundance (Fig. S5b) in LHCII trimers from active or inactive qH. Composition of the main chloroplastic lipids in LHCII trimer, LHCII/Lhcb monomer, or thylakoid extracts, indicated no significant differences (Fig. 4, S6); the distribution of thylakoid lipids is in line with the literature (Burgos et al., 2011). Of note, the 6-h cold and high light treatment did not alter the lipid profile significantly. The protein content was also similar in LHCII trimer from active or inactive qH (with an equivalent low amount of minor monomeric Lhcb4) and there were no visible additional protein bands or size shifts (Fig. S7). Investigation of possible post-translational modifications of amino acid residues by protein mass spectrometry will be the subject of future work. We observed LHCII subunits in the monomer fractions (probed with anti-Lhcb2), hence the “LHCII/Lhcb” denomination, with a higher content in the cold and high light treated samples compared to non-treated WT; this could be due to monomerization of trimers during the cold and high light treatment.

**Figure 4.**
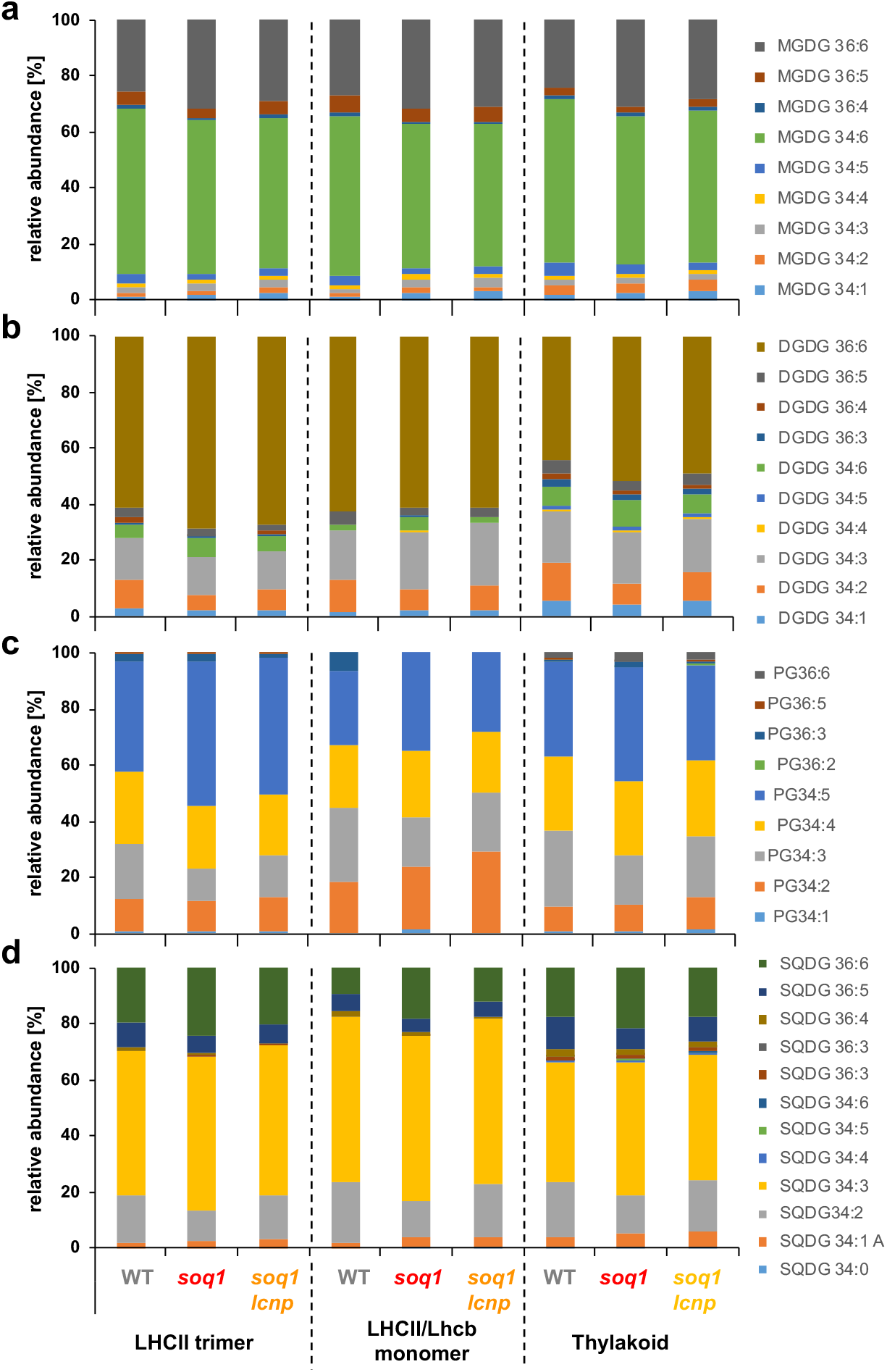
No significant differences in lipid composition with active or inactive qH. Lipid composition of pooled fractions from isolated LHCII trimer from non-treated wild type (WT) or cold HL-treated (replicate 3, n=8 plants) *soq1* (active qH) and *soq1 lcnp* (inactive qH) (see Fig. S3a, right panel for gel filtration experiment from which fractions were pooled). **(a)** monogalactosyldiacylglycerol (MGDG), **(b)** digalactosyldiacylglycerol (DGDG), **(c)** phosphatidylglycerol (PG) and **(d)** sulfoquinovosyldiacylglycerol (SQDG) species. Relative abundance is calculated per sample based on sum of given species present; absolute quantity normalized to chlorophyll content also did not show significant differences. Representative from two independent biological experiments is shown (see Fig. S6 for biological replicate 2).

### qH does not rely on a specific major LHCII subunit

Having gained the knowledge that qH occurs in the LHCII trimer, the next question was whether a specific LHCII subunit would be required, and this may provide a hint to the molecular origin of qH. We used genetic crosses together with genome editing to combine the *soq1* mutation with mutations in *LHCII* genes. The *soq1* mutant was crossed to *lhcb1* or *lhcb2* mutant lines generated by clustered regularly interspaced short palindromic repeat (CRISPR)-CRISPR-associated nuclease 9 (Cas9)-mediated genome editing or to the T-DNA insertional mutant *lhcb3*. The dissection of a putative specific LHCII quenching site is no small feat as there are five *LHCB1* genes (*LHCB1.1, 1.2*, and *1.3* are neighboring genes, so are *LHCB1.4* and *1.5*), three *LHCB2* genes (*LHCB2.1* and *2.2* are neighboring genes, and *LHCB2.3*) and one *LHCB3* gene. Three “loci” are therefore segregating upon generating the sextuple *soq1 lhcb1* or the quadruple *soq1 lhcb2* mutants. We genotyped the lines by PCR and confirmed lack of specific LHCII isoforms by immunoblot analysis (Fig. S8a). In all three mutant combinations, *soq1 lhcb1, soq1 lhcb2*, or *soq1 lhcb3*, additional quenching compared to the respective *lhcb* mutant controls was observed (Fig. 5a,c,e), which suggests that qH does not require a specific LHCII isoform; of note, NPQ can be compared between *lhcb* and *soq1 lhcb* mutants as they possess similar F_m_ values (Fig. 5b,d,f). In the case of *soq1 lhcb1*, only few trimers should be remaining (Pietrzykowska et al., 2014, Sattari Vayghan et al., 2022), but the NPQ difference between *lhcb1* and *soq1 lhcb1* is higher than between wild type and *soq1*. We therefore generated the *soq1 lhcb1 lcnp* to ensure that all additional quenching in *soq1 lhcb1* is qH (i.e., LCNP-dependent). The NPQ kinetics of *soq1 lhcb1 lcnp* and *lhcb1* were similar, which confirms that this additional quenching is qH and is enhanced when Lhcb1 is lacking (Fig.5a, S8b).

**Figure 5.**
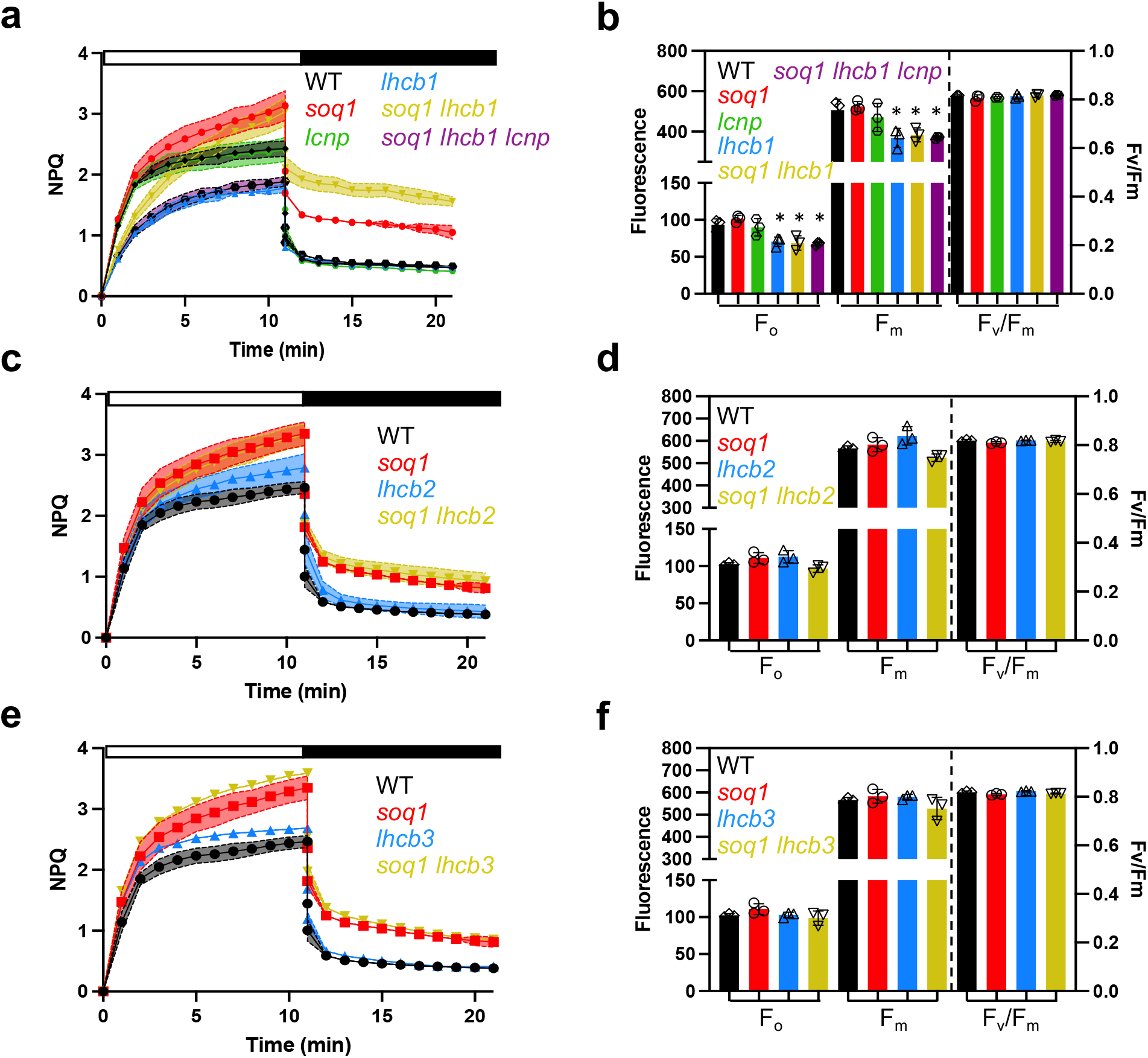
qH does not rely on a specific major Lhcb. **(a, c, e)** NPQ kinetics of WT, *soq1, lcnp, lhcb1, soq1 lhcb1, soq1 lhcb1 lcnp, lhcb2, soq1 lhcb2, lhcb3* and *soq1 lhcb3* four-week-old plants grown at 120 μmol photons m^-2^ s^-1^ dark-acclimated for 20 min. Induction of NPQ at 1300 μmol photons m^-2^ s^-1^ (white bar) and relaxation in the dark (black bar). **(b, d, f)** photosynthetic parameters F_o_, F_m_ and F_v_/F_m_ of the same plants. Tukey’s multiple comparisons test shows that *lhcb1, soq1 lhcb1* and *soq1 lhcb1 lcnp* are statistically different from WT for F_o_ (*P=0.0359, P*=0.0222 and *P*=0.0171, respectively) and F_m_ (*P*=0.0245, *P*=0.0482 and *P*=0.0257, respectively). Small significant difference in F_m_ with *P=0.0111* for *lhcb2* and *soq1 lhcb2* was not observed in two other biological experiments. Data represent means ± SD (*n*=3 plant individuals).

## Discussion

Here, we have characterized the chlorophyll fluorescence lifetimes of leaves, thylakoids, and isolated photosynthetic subcomplexes directly from *Arabidopsis* plants with active or inactive qH. We demonstrated that qH promotes a chlorophyll de-excitation pathway, which remains active upon isolation of thylakoid membranes (Fig. 1) and isolation of LHCII trimers (Fig. 2, 3) (see also summary in Table S2), but the effect is much smaller in isolated trimers than in leaves or thylakoids likely due to a large proportion of unquenched LHCII that dominates the ensemble.

The lifetime of non-treated *soq1 roqh1* leaves with constitutive activation of qH is among the shortest ever-observed with only ~0.1 ns (for comparison, a qE-induced state has a lifetime of ~0.5 ns (Sylak-Glassman et al., 2014) or the astaxanthin-synthesizing tobacco ~0.6 ns (Xu et al., 2020)) and confirms the stunted growth of *soq1 roqh1* (Amstutz et al., 2020) as being due to impaired light-harvesting function. Cold and high light-induced qH active plants display a short average fluorescence lifetime in leaves, similar to *soq1 roqh1* (~0.1 ns), and the lifetime increases in thylakoids (~0.4 ns) and is higher in isolated LHCII trimers (~2.6 ns). Whereas intact leaves and thylakoids of treated *soq1* with active qH showed a large amplitude of a rapidly decaying fluorescence component (A1>50%) and a small amplitude of a long-lived component (A3<10%), this trend was much less apparent at the level of isolated LHCII but yet remained true relative to *soq1 lcnp* (Table S3). Therefore, the long average fluorescence lifetime of LHCII isolated from treated *soq1* (~2.6 ns) is likely due to a decreased amplitude of rapid components (A1~18%) and increased amplitude of slow components (A3~73%) in the LHCII fraction relative to thylakoids. These results indicate that qH may relax during isolation of thylakoids or photosynthetic complexes (the latter takes about 8 h from leaf material collection to fluorescence lifetime measurements) and that the trimer fraction is a heterogenous population of quenched and unquenched LHCII. Furthermore, fluorescence lifetimes of pigment-protein complexes largely depend on their local environment, e.g. detergent or proteoliposome (Tietz et al., 2020, Crepin et al., 2021, Nicol & Croce, 2021), and comparison of LHCII in detergent micelles vs. membrane nanodiscs shows that quenching is attenuated by detergent (Son et al., 2020). Another possible explanation for the differences in fluorescence lifetimes among sample types is that a preserved membrane macroorganisation is required for a full qH response; indeed LH1 and LH2 antenna rings in purple bacteria display a 50% shorter lifetime *in vivo* compared to *in vitro* (Ricci et al., 1996), and similarly quenching in LHCII is dependent on its membrane environment (Moya et al., 2001, Natali et al., 2016, Saccon et al., 2020, Manna et al., 2021). In addition, other quenching sites beyond the LHCII trimers for qH may exist.

Nevertheless, the successful isolation of natively quenched LHCII by qH paves the way for revealing its molecular origin. We have not observed any significant changes in pigment, lipid or protein content of LHCII trimers with active qH (Fig. 4, Fig. S5-7). Because previous genetic dissection of qH requirement for xanthophyll pigments found that violaxanthin, zeaxanthin, or lutein are dispensable (Brooks et al., 2013, Malnoë et al., 2018), we tentatively propose that qH may stem from a chlorophyll-chlorophyll excitonic interaction state. Small changes in the conformation of the trimer modifying the protein environment of Chls or their orientation and/or distance with each other could enable qH, and identifying this fine-tuning will be the subject of future investigations. Such changes of the conformational space of proteins or carotenoids have recently been studied for qE experimentally or through molecular dynamics simulations (Liguori et al., 2017, Cignoni et al., 2021) and highlighted that several conformers would underlie light-harvesting and energy-dissipation states providing a more complex picture than previously thought for NPQ regulation.

Decreased fluorescence yield and lifetime of isolated LHCII trimer from plants with active qH indicates that qH likely occurs in the trimer (Fig. 2, 3). This interpretation of the results is assuming a similar relaxation rate between the different subcomplexes during isolation. The fractionation method used here results in a pool of LHCII trimers comprising the three types of trimers (strongly, moderately or loosely bound), so future work will tackle whether qH preferentially occurs in a specific type of trimer. Through genetic crosses, we found that qH does not require a specific LHCII subunit (Fig. 5), but it may rather require the trimeric conformation as the monomeric LHCII/Lhcb fraction displays similar fluorescence lifetimes whether qH is active or inactive (Fig. 3). LHCII trimers are composed of Lhcb1 (70% of the total LHCII proteins), Lhcb2 (20%) and Lhcb3 (10%) which form homo-trimers of Lhcb1, of Lhcb2, or hetero-trimers of Lhcb1, Lhcb2, and/or Lhcb3 (Standfuss & Kuhlbrandt, 2004). The degree of conservation between these subunits is high with an amino acid identity of 82% between Lhcb1 and Lhcb2, 78% between Lhcb1 and Lhcb3, and 79% between Lhcb2 and Lhcb3 (Caffarri et al., 2004). When a specific LHCII subunit is missing, some compensation by other subunits can occur: in the *lhcb1* CRISPR-Cas9 line, Lhcb2 accumulation is increased (Sattari Vayghan et al., 2022) but possibly insufficiently to fully explain the high NPQ of *soq1 lhcb1* (Fig. 5a). The enhanced qH in *soq1 lhcb1* could be explained by a different organization of photosynthetic complexes that would promote qH formation and/or slow down its relaxation. In the *amiLhcb2* or in *lhcb3* lines, trimers are abundant with an increased accumulation of Lhcb3 (Pietrzykowska et al., 2014), or Lhcb1 and 2 (Damkjaer et al., 2009) respectively. Therefore the similar NPQ kinetics between *soq1* and *soq1 lhcb2*, or *soq1 lhcb3*, and the enhanced qH in *soq1 lhcb1* (Fig. 5), indicate that qH does not rely on a specific subunit of the LHCII trimer.

To conclude, we isolated and characterized an energy-dissipative state of the major antenna complex directly from plants with active qH, with physiological relevance to natural conditions. Future work will focus on identifying differences in LHCII trimers that are associated with active qH and elucidation of the photophysical mechanism(s) of qH.

## Materials and Methods

### Plant material and growth conditions

Wild-type *Arabidopsis thaliana* and derived mutants studied here are of Col-0 ecotype. Mutants from these respective studies were used (only the *soq1-1* and *lcnp-1* alleles were used except for the *soq1 lhcb1 lcnp* line in which *lcnp* mutation was obtained through genome editing): *soq1* (Brooks et al., 2013), *soq1 lcnp* (Malnoë et al., 2018), *roqh1-1, −2, −3, soq1 roqh1-1, −2, −3, soq1 roqh1 ROQH1 OE, soq1 roqh1 lcnp, soq1 npq4 roqh1* (Amstutz et al., 2020), *lhcb3* (SALK_036200C) (Xu et al., 2012). For clarity, we refer to the *lhcb1* quintuple mutant affected in all five *LHCB1* genes as “*lhcb1*” (CRISPR-Cas9 edits for *lhcb1.1* (nucleotide insertion (nt) 575_576insA), *lhcb1.2* (nt deletion 575del), *lhcb1.3* (nt insertion 419_420insT), *lhcb1.4* (nt insertion 416_417insT), *lhcb1.5* (large deletion 413_581del)) and to the *lhcb2* triple mutant affected in all three *LHCB2* genes as “*lhcb2*” (*lhcb2.1* (SALK_005774C), CRISPR-Cas9 edits for *lhcb2.2* (nt insertion 10insA), *lhcb2.3* (nt insertion 93insA)). Mutants *soq1 lhcb1, soq1 lhcb2, soq1 lhcb3* and *soq1 lhcb1 lcnp* were generated in this study. Plants were grown on soil (Sunshine Mix 4/LA4 potting mix, Sun Gro Horticulture Distribution (Berkeley), 1:3 mixture of Agra-vermiculite “yrkeskvalité K-JORD” provided by RHP and Hasselfors garden respectively (Umeå)) under a 10/14 h light/dark photoperiod at 120 μmol photons m^-2^ s^-1^ at 21°C, referred to as standard conditions (Berkeley) or 8/16 h at 150 μmol photons m^-2^ s^-1^ at 22 °C/18°C (Umeå) for 5 to 6 weeks or seeds were surface sterilized using 70% ethanol and sown on agar plates (0.5 x Murashige and Skoog (MS) Basal Salt Mixture, Duchefa Biochemie, with pH adjusted to 5.7 with KOH) placed for 1 day in the dark at 4 °C, grown for 3 weeks with 12/12h at 150 μmol photons m^-2^ s^-1^ at 22°C and then transferred to soil. For the cold and high light treatment, plants or detached leaves were placed for 6 h at 6°C and at 1500 μmol photons m^-2^ s^-1^ using a custom-designed LED panel built by JBeamBio with cool white LEDs BXRA-56C1100-B-00 (Farnell). Light bulbs used in growth chambers are cool white (4100K) from Philips (F25T8/TL841 25W) for plants grown on soil and from General Electric (F17T8/SP41 17W) for seedlings grown on agar plates. A biological replicate, also referred to as biological experiment, represents a separate batch of several plant individuals grown at independent times. A technical replicate is an independent measurement performed on different aliquots from the same sample.

### Genetic crosses, genome editing and genotyping primers

Genetic crosses were done using standard techniques (Weigel & Glazebrook, 2006). Genome editing assisted by CRISPR-Cas9 was used to generate *lhcb1* and *lhcb2* and following the procedure described in (Ordon et al., 2017, Ordon et al., 2020) for the *soq1 lhcb1 lcnp* line. The mutant background *soq1 lhcb1* was used to generate the *soq1 lhcb1 lcnp* line using four sgRNA targeting *AtLCNP* exon 1 (CTTGTTGAAGTGGCAGCAGG), exon 3 (CTCACGTTACTGTCAGAAGA), exon 4 (TGACATCATAAGGCAACTTG) and exon 5 (TCAGTCACTTCACAGTCCTG) designed using the online tool CHOPCHOP (Labun et al., 2019) and further ranked for efficiency score with E-CRISP (Heigwer et al., 2014) for the two sgRNA targeting *AtLHCB1.1, LHCB1.2* and *LHCB1.3* CDS (GAGGACTTGCTTTACCCCGG) and *LHCB1.1, LHCB1.3, LHCB1.4* and *LHCB1.5* CDS (GGTTCACAGATCTTCAGCGA) and the two sgRNA targeting *LHCB2.2* exon 1 (GGATTGTTGGATAGCTGATG) and *LHCB2.3* exon 1 (GATGCGGCCACCGCCATTGG) in the background of a T-DNA insertional mutant for the *LHCB2.1* gene (SALK_005774C). The two sgRNAs targeting *LHCB1* or *LHCB2* genes were inserted into a binary vector under the control of the U6 promoter using the cloning strategy detailed by (Xing et al., 2014). This binary vector contains also the Cas9 gene under the control of the synthetic EC1 promoter that is expressed only in the egg cells (Durr et al., 2018). To identify *lhcb1* and *lhcb2*, resistant plants were screened by chlorophyll fluorescence for NPQ and photosynthetic acclimation based on (Pietrzykowska et al., 2014), and potential candidates were further confirmed by immunoblot using antibodies against Lhcb1 and Lhcb2. For *soq1 lhcb1 lcnp*, plants were transformed by floral dipping with Agrobacterium GV3101 pSoup containing the vector pDGE277 with the four sgRNAs. Seeds from transformed plants were plated and selected on MS plates with 25 μg mL^-1^ hygromycin. The hygromycin-resistant plants were selected and the absence of LCNP was confirmed by immunoblot using an antibody raised against LCNP. Phire Plant Direct PCR kit was used for genotyping and sequencing with dilution protocol (ThermoFisher Scientific F130); primer list can be found in Table S4.

### Chlorophyll fluorescence imaging

Chlorophyll fluorescence was measured at room temperature with Walz Imaging-PAM Maxi (Fig. S3b,c) or with SpeedZenII from JBeamBio (Fig. 5, S4c). For NPQ measurements, plants or detached leaves were dark-acclimated for 20 min and NPQ was induced by 1200 μmol photons m^-2^ s^-1^ for 10 min and relaxed in the dark for 10 min. Maximum fluorescence after dark acclimation (F_m_) and throughout measurement (F_m_’) were recorded after applying a saturating pulse of light. NPQ was calculated as (F_m_-F_m_’)/Fm’. F_v_/F_m_ is calculated as (F_m_-F_o_)/F_m_ where F_o_ is the minimum fluorescence after dark acclimation.

### Thylakoid extraction

Thylakoid extractions were performed according to (Iwai et al., 2015). Briefly, leaves from 6 to 8-week-old plants were ground in a blender for 30 s in 60 mL B1 cold solution (20 mM Tricine-KOH pH7.8, 400 mM NaCl, 2 mM MgCl_2_). Protease inhibitors are used at all steps (0.2 mM benzamidine, 1 mM aminocaproic acid, 0.2 mM PMSF). The solution is then filtrated through four layers of Miracloth and centrifuged 5 minutes at 27,000 x g at 4°C. The supernatant is discarded, and the pellet is resuspended in 15 mL B2 solution (20 mM Tricine-KOH pH 7.8, 150 mM NaCl, 5 mM MgCl_2_). The resuspended solution is overlayed onto a 1.3 M/1.8 M sucrose cushion and ultracentrifuged for 30 min in a SW28 rotor at 131,500 x g and 4°C. The band between the sucrose layers is removed and washed with B3 solution (20 mM Tricine-KOH pH 7.8, 15 mM NaCl, 5 mM MgCl_2_). The solution is centrifuged for 15 min at 27,000 x g and 4°C. The pellet is washed in storing solution (20 mM Tricine-KOH pH 7.8, 0.4 M sucrose, 15 mM NaCl, 5 mM MgCl_2_) and centrifuged for 10 min at 27,000 x g and 4°C. The pellet is then resuspended in storing solution. Chl concentration is measured according to (Porra et al., 1989). If samples are to be stored, they were flash-frozen in liquid nitrogen and stored at −80°C at approximately 2.5 mg Chl mL^-1^. Upon using thylakoid preparation, samples are rapidly thawed and buffer is exchanged with 120 mM Tris-HCl pH 6.8, and Chl concentration is measured. For spectroscopy experiments, thylakoids were isolated according to (Gilmore et al., 1998). For the ‘non-treated’ condition, leaves were detached and dark-acclimated overnight at 4°C.

### Isolation of pigment-protein complexes

Thylakoids membranes (400 μg Chl) were solubilized at 2 mg mL^-1^ with 4% (w/v) α-dodecyl maltoside (α-DM) for 15 min on ice (solution was briefly mixed every 5 min), unsolubilized membranes were removed by centrifugation at 14,000 rpm for 5 min. Gel filtration chromatography was performed as described in (Iwai et al., 2015) using the ÄKTAmicro chromatography system with a Superdex 200 Increase 10/300 GL column (GE Healthcare) equilibrated with 20 mM Tris-HCl (pH 8.0), 5 mM MgCl_2_ and 0.03% (w/v) α-DM at room temperature. The flow rate was 1 mL min^-1^. The proteins were detected at 280 nm absorbance.

### Protein analysis

A 5 mm diameter disc was cut from the leaf and frozen into liquid nitrogen. The leaf disc was ground with a plastic pestle and 100 μL of sample loading buffer (62.5 mM Tris pH7.5, 2% SDS, 10% Glycerol, 0.01% Bromophenol blue, 100 mM DTT) was added. Samples were boiled at 95-100°C for 5 min and centrifuged for 3 min. From the samples, 10 μL were loaded onto a 10% SDS-PAGE gel. For the gel filtration fractions, samples were loaded at same volume from pooled adjacent fractions (three fractions for each) onto a 12% SDS-PAGE gel for immunoblot or for silver stain. After migration the proteins were transferred to a PVDF 0.45 μm from ThermoScientific. After transferring the membrane was blocked with TBST+ 3% milk for 1h followed by 1h incubation of the primary antibody (ATP*β* AS05 085 (1:5,000), Lhcb1 AS09 522 (1:5,000), Lhcb2 AS01 003 (1:10,000), Lhcb3 AS01 002 (1:2,000), Lhcb4 AS04 045 (1:7,000) from Agrisera and rabbit antibodies against a peptide of LCNP (AEDLEKSETDLEKQ) were produced and purified by peptide affinity by Biogenes and used at a 1:200 dilution) diluted in TBST+ 3% milk. The membrane was washed 3 times 10 min with TBST. Then incubated for 1h with the secondary goat anti-rabbit antibody conjugated to horse radish peroxidase AS09 602 (1:10,000) from Agrisera in TBST + 3% milk. The membrane was washed 3 times 10 min with TBST and 1 time 5 min with TBS. The Agrisera ECL Bright (AS16 ECL-N-100) and Azure biosystems c600 were used to reveal the bands.

### Clear-native PAGE analysis

Thylakoid are washed with the solubilization buffer (25 mM BisTris/HCl (pH 7.0), 20% (w/v) glycerol, 1mM *ε*-aminocaproic acid and 0.2 mM PMSF) and resuspended in the same buffer at 1 mg Chl mL^-1^. An equal volume of 2% α-DM was added to the thylakoid solution for 15 min on ice in the dark. Traces of insoluble material were removed by centrifugation at 18,000 x g 20 min 4°C. The Chl concentration was measured, and proteins were loaded at equal Chl content in the native gel (NativePAGE™ 3 to 12%, Bis-Tris, 1.0 mm, Mini Protein Gel, 10-well from ThermoFisher catalog number BN1001BOX). Prior to loading the samples were supplemented with sodium deoxycholate (final concentration 0.3%). The cathode buffer is 50 mM Tricine, 15 mM BisTris, 0.05% sodium deoxycholate and 0.02% α-DM, pH7.0 and anode buffer is 50 mM BisTris, pH7.0. Electrophoresis was performed at 4°C with a gradual increase in voltage: 75 V for 30 min, 100 V for 30 min, 125 V for 30 min, 150 V for 1 h and 175 V until the sample reached the end of the gel. Method adapted from (Rantala et al., 2018).

### Pigment extraction and analysis

HPLC analysis of carotenoids and Chls was done as previously described (Müller-Moulé et al., 2002). 10 μg Chl of fraction samples were extracted in 200 μl 100% acetone.

### Lipid profiling

Thylakoids or gel filtration fractions corresponding to trimers or monomers were evaporated until dryness using a vacuum evaporator and dried samples were reconstituted in 100 μl isopropanol. Lipids were separated on Acquity Ultra Performance LC coupled to a Synapt G2 HDMS equipped with electrospray ionization source (Waters) according to an adapted protocol (Mueller et al., 2015). Briefly, liquid chromatographic separation was performed on BEH C18 column (2.1×100 mm, 1.7 μm) using binary solvent strength gradient from 30% to 100% eluent B within 10 minutes at a flow rate of 0.3 mL min^-1^. Eluent A was 10 mM ammonium acetate in water:acetonitrile (60:40 v/v) and eluent B was 10 mM ammonium acetate in isopropanol:acetonitrile (90:10 v/v). The mass spectrometer was operated in positive and negative electrospray ionisation and centroid data was acquired with a mass range from 50-1200 Da using leucine-enkephaline for internal calibration. Lipids were identified by matching masses of molecular, typical fragments (error less than 1 mDa) and elemental compositions using isotope abundance distributions. MassLynx 4.1. was used to operate the instrument and QuantLynx was used for peak integration. Samples were normalized by chlorophyll content.

### Fluorescence spectroscopy on isolated thylakoids or complexes

Room temperature fluorescence emission of gel filtration fractions and dependence on step solubilization of thylakoids were performed according to (Dall’Osto et al., 2005) using a Horiba FluoroMax fluorimeter and Starna cells 18/9F-SOG-10 (path length 10mm) with Chl concentration of 0.1 μg mL^-1^. For the emission spectrum of gel filtration fractions (emission 650 to 800 nm with excitation at 625 nm, bandwidth, 5 nm for excitation, 5 nm for emission), samples were diluted at same absorption (Δ625-750 nm=0.0005) in 20 mM Tris HCl pH8, 5 mM MgCl_2_ and 0.03% α-DM. For the step solubilization (emission 680 nm with excitation at 440 nm, bandwidth, 5 nm for excitation, 3 nm for emission), thylakoids preparation were diluted in 20 mM Tris HCl pH8, 5 mM MgCl_2_, and two different detergents were added: first, α-DM at final 0.5% (w/v) concentration from a 10% stock solution which dissociates the pigment binding proteins from each other without release of chlorophyll from their protein moiety (Caffarri et al., 2001), then Triton X-100 at final 5% (w/v) concentration from a 50% stock solution which denatures the pigment–proteins and yields free pigments (Giuffra et al., 1997). After each addition, the cuvette was turned upside-down 3 to 5 times for mixing and time for fluorescence level stabilization was allowed.

### Fluorescence lifetime measurements and fitting

Method used is adapted for fluorescence lifetime snapshot from (Sylak-Glassman et al., 2016). Time-correlated single photon counting (TCSPC) was performed on detached leaves, isolated thylakoids and gel filtration fractions. Excitation at 420 nm was provided by frequency doubling the 840 nm output of a Ti:sapphire oscillator (Coherent, Mira 900f, 76 MHz). The laser intensity was ~18,000 μmol photons m^-2^ s^-1^/pulse (~20 pJ/pulse), sufficient to close reaction centers. Emission was monitored at 680 nm using a MCP PMT detector (Hamamatsu, R3809U). The FWHM of IRF was ~30-40 ps.

It has been shown that a wide range of exponentials can reasonably fit any ensemble fluorescence decay measurement (Bennett et al., 2018), with no easy way to distinguish between the different “models”. Therefore to gain a simple, unbiased description of the fluorescence dynamics in each sample, each decay was fit to a tri-exponential model (Picoquant, Fluofit Pro-4.6) without constraining any specific kinetic component, and an amplitude-weighted average fluorescence lifetime (τ_avg_) was calculated. The extent of quenching was then evaluated by comparison of τ_avg_ values from non-treated and cold HL-treated plants, quantified as 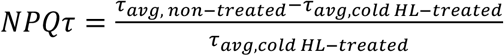. Prior to each measurement, qE was relaxed by dark-acclimation for at least 5 min.

## Author Contributions

A.M., K.K.N. and G.R.F. designed research; A.M., S.P., C.J.S., E.J.S.-G., C.L.A. and L.L. performed biochemical isolation and spectroscopy and analyzed the data; A.F. performed lipid profiling and A.F. and M.J.M. analyzed the data; F.L. generated the *lhcb1* and *lhcb2* lines; P.B. performed crosses, genome editing, native gels and analyses of *lhcb* mutant combinations. All of the authors discussed the data, and A.M. wrote the paper with input from P.B., C.J.S., G.R.F and K.K.N.

## Acknowledgments

We thank Masakazu Iwai for advice and technical assistance regarding isolation of pigment-protein complexes and for critical reading of the manuscript, Alexandra Lee Fisher for helpful discussions regarding fluorescence lifetimes experiments, Maria Lesch for assistance with lipid profiling, Julie Guerreiro for assistance with antenna mutant combination analysis, Jingfang Hao for generation of a new LCNP antibody and Yolande Provot for assistance with isolation of the *soq1 lhcb1 lcnp* line; Roberta Croce, Roberto Bassi for critical discussions. This work used the Metabolomics Core Unit of the University Wuerzburg, supported by the German Research Foundation Deutsche Forschungsgemeinschaft (project 179877739), for lipid profiling. This research (Berkeley) was supported by the Division of Chemical Sciences, Geosciences and Biosciences, Office of Basic Energy Sciences, Office of Science, US Department of Energy (Field Work Proposal 449B). K.K.N. is an investigator of the Howard Hughes Medical Institute. A.M. (Umeå) was supported by European Commission Marie Skłodowska-Curie Actions Individual Fellowship Reintegration Panel (845687). The research performed at Umeå (P.B. and A.M.) was supported by grants to UPSC from the Knut and Alice Wallenberg Foundation (2016.0341 and 2016.0352), the Swedish Governmental Agency for Innovation Systems (2016-00504), by a starting grant to A.M. from the Swedish Research Council Vetenskapsrådet (2018-04150) and by a consortium grant from the Swedish Foundation for Strategic Research (ARC19-0051). The research performed at Neuchâtel (F.L.) was supported by the Swiss National Science Foundation (31003A_179417).

## Data availability

The authors declare that all data supporting the findings of this study are included in the paper and its supplementary information file, and are available from the corresponding author upon request. Source data for all figures are provided with the paper. Sequence data from this article can be found in the Arabidopsis Genome Initiative under accession numbers At1g29920 (Lhcb1.1), At1g29910 (Lhcb1.2), At1g29930 (Lhcb1.3), At1g56500 (SOQ1), At2g05070 (Lhcb2.2), At2g05100 (Lhcb2.1), At2g34420 (Lhcb1.5), At2g34430 (Lhcb1.4), At3g27690 (Lhcb2.3), At3g47860 (LCNP), At4g31530 (ROQH1), At5g54270 (Lhcb3).

## Supplemental Data

Fig.S1 to S8 and Table S1 to S4.

## References

Amstutz CL, Fristedt R, Schultink A, Merchant SS, Niyogi KK, Malnoë A (2020) An atypical short-chain dehydrogenase-reductase functions in the relaxation of photoprotective qH in Arabidopsis. Nat Plants 6: 154–166

Ballottari M, Girardon J, Dall’osto L, Bassi R (2012) Evolution and functional properties of photosystem II light harvesting complexes in eukaryotes. Biochim Biophys Acta 1817: 143–57

Bassi R, Dall’Osto L (2021) Dissipation of Light Energy Absorbed in Excess: The Molecular Mechanisms. Annu Rev Plant Biol 72: 47–76

Bennett DIG, Fleming GR, Amarnath K (2018) Energy-dependent quenching adjusts the excitation diffusion length to regulate photosynthetic light harvesting. Proc Natl Acad Sci U S A 115: E9523–E9531

Brooks MD, Sylak-Glassman EJ, Fleming GR, Niyogi KK (2013) A thioredoxin-like/beta-propeller protein maintains the efficiency of light harvesting in Arabidopsis. Proc Natl Acad Sci USA 110: 2733–40

Bru P, Nanda S, Malnoë A (2020) A Genetic Screen to Identify New Molecular Players Involved in Photoprotection qH in Arabidopsis thaliana. Plants 9: 1565

Burgos A, Szymanski J, Seiwert B, Degenkolbe T, Hannah MA, Giavalisco P, Willmitzer L (2011) Analysis of short-term changes in the Arabidopsis thaliana glycerolipidome in response to temperature and light. Plant J 66: 656–68

Caffarri S, Croce R, Breton J, Bassi R (2001) The major antenna complex of photosystem II has a xanthophyll binding site not involved in light harvesting. J Biol Chem 276: 35924–33

Caffarri S, Croce R, Cattivelli L, Bassi R (2004) A look within LHCII: differential analysis of the Lhcb1-3 complexes building the major trimeric antenna complex of higher-plant photosynthesis. Biochemistry 43: 9467–76

Cignoni E, Lapillo M, Cupellini L, Acosta-Gutierrez S, Gervasio FL, Mennucci B (2021) A different perspective for nonphotochemical quenching in plant antenna complexes. Nat Commun 12: 7152

Crepin A, Caffarri S (2018) Functions and Evolution of Lhcb Isoforms Composing LHCII, the Major Light Harvesting Complex of Photosystem II of Green Eukaryotic Organisms. Curr Protein Pept Sci 19: 699–713

Crepin A, Cunill-Semanat E, Kuthanova Trskova E, Belgio E, Kana R (2021) Antenna Protein Clustering In Vitro Unveiled by Fluorescence Correlation Spectroscopy. Int J Mol Sci 22

Dall’Osto L, Caffarri S, Bassi R (2005) A mechanism of nonphotochemical energy dissipation, independent from PsbS, revealed by a conformational change in the antenna protein CP26. Plant Cell 17: 1217–32

Damkjaer JT, Kereiche S, Johnson MP, Kovacs L, Kiss AZ, Boekema EJ, Ruban AV, Horton P, Jansson S (2009) The photosystem II light-harvesting protein Lhcb3 affects the macrostructure of photosystem II and the rate of state transitions in Arabidopsis. Plant Cell 21: 3245–56

Demmig-Adams B, Garab G, Adams III W, Govindjee UoI (2014) Non-photochemical quenching and energy dissipation in plants, algae and cyanobacteria. In Springer

Durr J, Papareddy R, Nakajima K, Gutierrez-Marcos J (2018) Highly efficient heritable targeted deletions of gene clusters and non-coding regulatory regions in Arabidopsis using CRISPR/Cas9. Sci Rep 8: 4443

Gilmore AM, Shinkarev VP, Hazlett TL, Govindjee G (1998) Quantitative analysis of the effects of intrathylakoid pH and xanthophyll cycle pigments on chlorophyll a fluorescence lifetime distributions and intensity in thylakoids. Biochemistry 37: 13582–93

Giuffra E, Zucchelli G, Sandona D, Croce R, Cugini D, Garlaschi FM, Bassi R, Jennings RC (1997) Analysis of some optical properties of a native and reconstituted photosystem II antenna complex, CP29: pigment binding sites can be occupied by chlorophyll a or chlorophyll b and determine spectral forms. Biochemistry 36: 12984–93

Heigwer F, Kerr G, Boutros M (2014) E-CRISP: fast CRISPR target site identification. Nat Methods 11: 122–3

Horton P, Ruban AV, Walters RG (1996) Regulation of Light Harvesting in Green Plants. Annu Rev Plant Physiol Plant Mol Biol 47: 655–684

Ilioaia C, Johnson MP, Horton P, Ruban AV (2008) Induction of efficient energy dissipation in the isolated light-harvesting complex of Photosystem II in the absence of protein aggregation. J Biol Chem 283: 29505–12

Iwai M, Yokono M, Kono M, Noguchi K, Akimoto S, Nakano A (2015) Light-harvesting complex Lhcb9 confers a green alga-type photosystem I supercomplex to the moss Physcomitrella patens. Nat Plants 1: 14008

Jahns P, Holzwarth AR (2012) The role of the xanthophyll cycle and of lutein in photoprotection of photosystem II. Biochim Biophys Acta 1817: 182–93

Krause GH, Somersalo S, Zumbusch E, Weyers B, Laasch H (1990) On the Mechanism of Photoinhibition in Chloroplasts. Relationship Between Changes in Fluorescence and Activity of Photosystem II. Journal of Plant Physiology 136: 472–479

Kromdijk J, Glowacka K, Leonelli L, Gabilly ST, Iwai M, Niyogi KK, Long SP (2016) Improving photosynthesis and crop productivity by accelerating recovery from photoprotection. Science 354: 857–861

Labun K, Montague TG, Krause M, Torres Cleuren YN, Tjeldnes H, Valen E (2019) CHOPCHOP v3: expanding the CRISPR web toolbox beyond genome editing. Nucleic Acids Res 47: W171–W174

Levesque-Tremblay G, Havaux M, Ouellet F (2009) The chloroplastic lipocalin AtCHL prevents lipid peroxidation and protects Arabidopsis against oxidative stress. Plant J 60: 691–702

Liguori N, Periole X, Marrink SJ, Croce R (2015) From light-harvesting to photoprotection: structural basis of the dynamic switch of the major antenna complex of plants (LHCII). Sci Rep 5: 15661

Liguori N, Xu P, van Stokkum IHM, van Oort B, Lu Y, Karcher D, Bock R, Croce R (2017) Different carotenoid conformations have distinct functions in light-harvesting regulation in plants. Nat Commun 8: 1994

Malnoё A (2018) Photoinhibition or photoprotection of photosynthesis? Update on the (newly termed) sustained quenching component, qH. Environmental and Experimental Botany 154: 123–133

Malnoё A, Schultink A, Shahrasbi S, Rumeau D, Havaux M, Niyogi KK (2018) The Plastid Lipocalin LCNP Is Required for Sustained Photoprotective Energy Dissipation in Arabidopsis. Plant Cell 30: 196–208

Manna P, Davies T, Hoffmann M, Johnson MP, Schlau-Cohen GS (2021) Membrane-dependent heterogeneity of LHCII characterized using single-molecule spectroscopy. Biophysical Journal 120: 3091–3102

Moya I, Silvestri M, Vallon O, Cinque G, Bassi R (2001) Time-Resolved Fluorescence Analysis of the Photosystem II Antenna Proteins in Detergent Micelles and Liposomes. Biochemistry 40: 12552–12561

Mueller SP, Krause DM, Mueller MJ, Fekete A (2015) Accumulation of extra-chloroplastic triacylglycerols in Arabidopsis seedlings during heat acclimation. Journal of Experimental Botany 66: 4517–4526

Müller P, Li XP, Niyogi KK (2001) Non-photochemical quenching. A response to excess light energy. Plant Physiol 125: 1558–66

Müller-Moulé P, Conklin PL, Niyogi KK (2002) Ascorbate deficiency can limit violaxanthin deepoxidase activity in vivo. Plant Physiol 128: 970–7

Natali A, Gruber JM, Dietzel L, Stuart MCA, van Grondelle R, Croce R (2016) Lightharvesting Complexes (LHCs) Cluster Spontaneously in Membrane Environment Leading to Shortening of Their Excited State Lifetimes*. Journal of Biological Chemistry 291: 16730–16739

Nawrocki WJ, Liu X, Raber B, Hu C, de Vitry C, Bennett DIG, Croce R (2021) Molecular origins of induction and loss of photoinhibition-related energy dissipation qI. Sci Adv 7: eabj0055

Nicol L, Croce R (2021) The PsbS protein and low pH are necessary and sufficient to induce quenching in the light-harvesting complex of plants LHCII. Sci Rep 11: 7415

Niyogi KK, Truong TB (2013) Evolution of flexible non-photochemical quenching mechanisms that regulate light harvesting in oxygenic photosynthesis. Curr Opin Plant Biol 16: 307–14

Ordon J, Bressan M, Kretschmer C, Dall’Osto L, Marillonnet S, Bassi R, Stuttmann J (2020) Optimized Cas9 expression systems for highly efficient Arabidopsis genome editing facilitate isolation of complex alleles in a single generation. Funct Integr Genomics 20: 151–162

Ordon J, Gantner J, Kemna J, Schwalgun L, Reschke M, Streubel J, Boch J, Stuttmann J (2017) Generation of chromosomal deletions in dicotyledonous plants employing a user-friendly genome editing toolkit. Plant J 89: 155–168

Ort DR, Merchant SS, Alric J, Barkan A, Blankenship RE, Bock R, Croce R, Hanson MR, Hibberd JM, Long SP, Moore TA, Moroney J, Niyogi KK, Parry MA, Peralta-Yahya PP, Prince RC, Redding KE, Spalding MH, van Wijk KJ, Vermaas WF et al. (2015) Redesigning photosynthesis to sustainably meet global food and bioenergy demand. Proc Natl Acad Sci U S A 112: 8529–36

Pietrzykowska M, Suorsa M, Semchonok DA, Tikkanen M, Boekema EJ, Aro EM, Jansson S (2014) The light-harvesting chlorophyll a/b binding proteins Lhcb1 and Lhcb2 play complementary roles during state transitions in Arabidopsis. Plant Cell 26: 3646–60

Pinnola A, Bassi R (2018) Molecular mechanisms involved in plant photoprotection. Biochem Soc Trans 46: 467–482

Porra RJ, Thompson WA, Kriedemann PE (1989) Determination of accurate extinction coefficients and simultaneous equations for assaying chlorophylls a and b extracted with four different solvents: verification of the concentration of chlorophyll standards by atomic absorption spectroscopy. Biochimica et Biophysica Acta (BBA) - Bioenergetics 975: 384–394

Quick WP, Stitt M (1989) An examination of factors contributing to non-photochemical quenching of chlorophyll fluorescence in barley leaves. Biochim Biophys Acta 977: 287–296

Rantala M, Paakkarinen V, Aro E-M (2018) Analysis of Thylakoid Membrane Protein Complexes by Blue Native Gel Electrophoresis. JoVE: e58369

Ricci M, Bradforth SE, Jimenez R, Fleming GR (1996) Internal conversion and energy transfer dynamics of spheroidene in solution and in the LH-1 and LH-2 light-harvesting complexes. Chemical Physics Letters 259: 381–390

Saccon F, Durchan M, Bína D, Duffy CDP, Ruban AV, Polívka T (2020) A Protein Environment-Modulated Energy Dissipation Channel in LHCII Antenna Complex. iScience 23

Sattari Vayghan H, Nawrocki WJ, Schiphorst C, Tolleter D, Hu C, Douet V, Glauser G, Finazzi G, Croce R, Wientjes E, Longoni F (2022) Photosynthetic Light Harvesting and Thylakoid Organization in a CRISPR/Cas9 Arabidopsis Thaliana LHCB1 Knockout Mutant. Frontiers in Plant Science 13

Son M, Pinnola A, Gordon SC, Bassi R, Schlau-Cohen GS (2020) Observation of dissipative chlorophyll-to-carotenoid energy transfer in light-harvesting complex II in membrane nanodiscs. Nat Commun 11: 1295

Standfuss J, Kuhlbrandt W (2004) The three isoforms of the light-harvesting complex II: spectroscopic features, trimer formation, and functional roles. J Biol Chem 279: 36884–91

Sylak-Glassman EJ, Malnoë A, De Re E, Brooks MD, Fischer AL, Niyogi KK, Fleming GR (2014) Distinct roles of the photosystem II protein PsbS and zeaxanthin in the regulation of light harvesting in plants revealed by fluorescence lifetime snapshots. Proc Natl Acad Sci USA 111: 17498–503

Sylak-Glassman EJ, Zaks J, Amarnath K, Leuenberger M, Fleming GR (2016) Characterizing non-photochemical quenching in leaves through fluorescence lifetime snapshots. Photosynth Res 127: 69–76

Takabayashi A, Kurihara K, Kuwano M, Kasahara Y, Tanaka R, Tanaka A (2011) The Oligomeric States of the Photosystems and the Light-Harvesting Complexes in the Chl *b*-Less Mutant. Plant and Cell Physiology 52: 2103–2114

Tietz S, Leuenberger M, Hohner R, Olson AH, Fleming GR, Kirchhoff H (2020) A proteoliposome-based system reveals how lipids control photosynthetic light harvesting. J Biol Chem 295: 1857–1866

Valkunas L, Chmeliov J, Krüger TPJ, Ilioaia C, van Grondelle R (2012) How Photosynthetic Proteins Switch. The Journal of Physical Chemistry Letters 3: 2779–2784

van Oort B, van Hoek A, Ruban AV, van Amerongen H (2007) Equilibrium between quenched and nonquenched conformations of the major plant light-harvesting complex studied with high-pressure time-resolved fluorescence. J Phys Chem B 111: 7631–7

Wang Y, Burgess SJ, de Becker EM, Long SP (2020) Photosynthesis in the fleeting shadows: an overlooked opportunity for increasing crop productivity? Plant J 101: 874–884

Weigel D, Glazebrook J (2006) Setting Up Arabidopsis Crosses. Cold Spring Harbor Protocols

Xing HL, Dong L, Wang ZP, Zhang HY, Han CY, Liu B, Wang XC, Chen QJ (2014) A CRISPR/Cas9 toolkit for multiplex genome editing in plants. BMC Plant Biol 14: 327

Xu P, Chukhutsina VU, Nawrocki WJ, Schansker G, Bielczynski LW, Lu Y, Karcher D, Bock R, Croce R (2020) Photosynthesis without β-carotene. eLife 9: e58984

Xu Y-H, Liu R, Yan L, Liu Z-Q, Jiang S-C, Shen Y-Y, Wang X-F, Zhang D-P (2012) Lightharvesting chlorophyll a/b-binding proteins are required for stomatal response to abscisic acid in Arabidopsis. Journal of experimental botany 63: 1095–1106

Zhu XG, Long SP, Ort DR (2010) Improving photosynthetic efficiency for greater yield. Annu Rev Plant Biol 61: 235–61

