## Supplemental Data for "An energy-dissipative state of the major antenna complex of plants"

**Supplemental Data:** Fig. S1 to S8 and Table S1 to S4

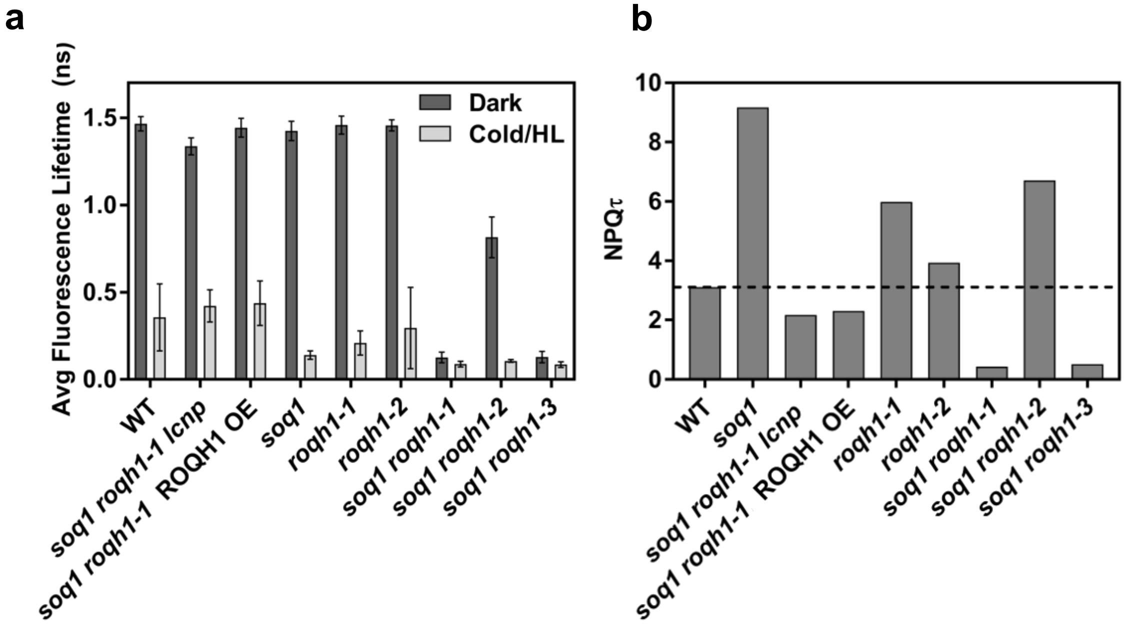


**Fig. S1. Fluorescence lifetime from leaves with active qH are shorter (supports Fig. 1).**
**(a)** Average fluorescence lifetimes determined by time-correlated single photon counting from non-treated plants (dark) and from cold and high light-treated plants for 6 h at 6°C and 1500 µmol photons $m^{-2}$ $s^{-1}$ (cold HL). Plants were dark acclimated for 10 min prior to measurement on individual leaves. Data represent mean ± SD, n = 20-24 measurements on n = 10-12 leaves taken from 2-3 plants. Each leaf was measured once and then repositioned for a second laser measurement on a distinct region of the same leaf. **(b)** NPQτ calculated as (τdark – τcold HL) / τcold HL.

**Table S1. Fluorescence lifetime from leaves with active qH are shorter (supports Fig. 1).** Data displayed in Fig. S1 (ROQH1 OE is *soq1 roqh1-1* ROQH1 overexpressor) and data on WT and *soq1 npq4 roqh1* from non-treated plants (dark) and after a 10-min high light (HL) exposure at 1200 µmol photons $m^{-2}$ $s^{-1}$.

| Genotype | Avg Dark Lifetime (ns) ± SD | Avg Cold HL Lifetime (ns) ± SD | Calculated NPQ$_T$ |
|---|---|---|---|
| **WT** | 1.465 ± 0.041 | 0.356 ± 0.192 | 3.11 |
| ***roqh1-1*** | 1.458 ± 0.052 | 0.209 ± 0.069 | 5.99 |
| ***roqh1-2*** | 1.456 ± 0.032 | 0.295 ± 0.233 | 3.94 |
| ***roqh1-3*** | N/A (bolted) | N/A (bolted) | N/A (bolted) |
| **ROQH1 OE** | 1.443 ± 0.054 | 0.437 ± 0.127 | 2.30 |
| ***soq1*** | 1.424 ± 0.055 | 0.140 ± 0.024 | 9.17 |
| ***soq1 roqh1-1*** | 0.126 ± 0.031 | 0.088 ± 0.016 | 0.43 |
| ***soq1 roqh1-2*** | 0.815 ± 0.117 | 0.106 ± 0.008 | 6.71 |
| ***soq1 roqh1-3*** | 0.128 ± 0.033 | 0.085 ± 0.016 | 0.51 |
| ***soq1 roqh1 lcnp*** | 1.336 ± 0.049 | 0.422 ± 0.092 | 2.17 |

| Genotype | Avg Dark Lifetime (ns) ± SD | Avg 10-min HL Lifetime (ns) ± SD |
|---|---|---|
| **WT** | 1.27 ± 0.08 | 0.36 ± 0.02 |
| ***soq1 npq4 roqh1-1*** | 0.056 ± 0.002 | 0.053 ± 0.002 |

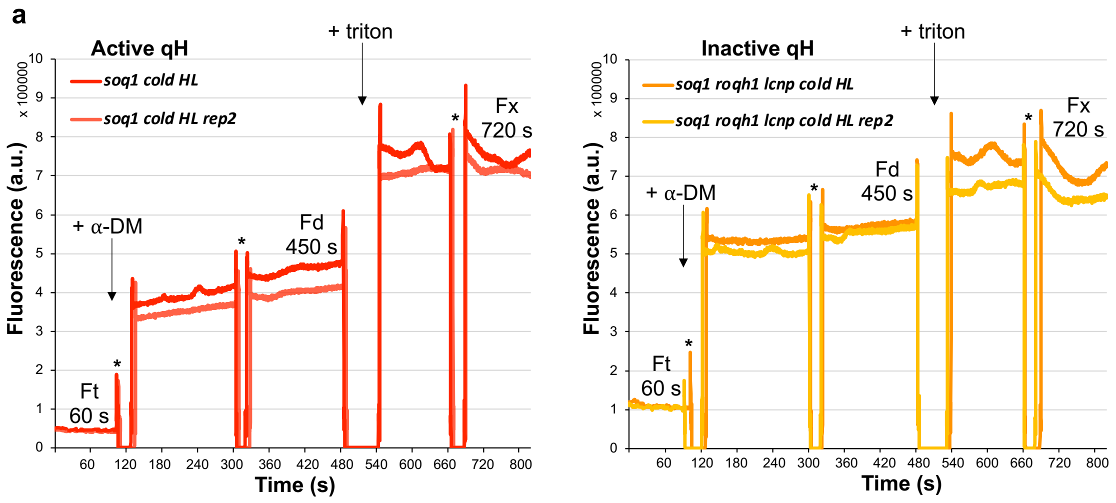


**b**

| | *soq1* non-treated | *soq1 roqh1 lcnp* non-treated | WT cold HL | *soq1 roqh1 lcnp* cold HL | *soq1* cold HL |
|---|---|---|---|---|---|
| Qm | 3.0 | 2.6 | 4.5 | 4.4 | **9.2** |
| Qpi | 0.26 | 0.24 | 0.3 | 0.2 | **0.7** |
| Qm+pi | 4 | 3.4 | 6.2 | 5.4 | **16.2** |

**Fig. S2. qH is partly provided by incorporation of chlorophyll-protein complexes in the thylakoid membrane and by chlorophyll binding into complexes. (a)** Release of chlorophyll fluorescence quenching by step solubilization of thylakoid membranes from cold and high light-treated *soq1*, active qH, and *soq1 roqh1 lcnp*, inactive qH. Two technical replicates (rep) are shown from one biological experiment with n = 8 plants. Fluorescence emission at 680 nm at room temperature in arbitrary units (a.u.) with excitation at 440 nm vs. time (s); samples were at same chlorophyll concentration (0.1 µg $mL^{-1}$) and were homogenized by taking out cuvette and turning it upside-down 3 to 5 times at symbol (*). Ft, fluorescence from unsolubilized membrane. Addition of α-DM 0.5% at 100 s for solubilization of the membranes which dissociates the pigment binding proteins from each other without release of chlorophyll from their protein moiety. Fd, fluorescence from pigment-protein complexes in α-DM. Addition of triton X-100 5% denatures the pigment–proteins and yields free pigments, generating the maximum quantum yield of fluorescence obtainable for the system (Fx). **(b)** Calculated quenching values of non-treated *soq1* and *soq1 roqh1 lcnp* (one technical replicate), and cold HL-treated WT, *soq1* and *soq1 roqh1 lcnp* (means of two technical replicates). Qm=(Fd-Ft)/Ft, quenching of chlorophyll–protein complexes fluorescence provided by their incorporation in the thylakoid lipid bilayer (due to protein–protein and lipid–proteins interactions in the membrane); Qpi=(Fx-Fd)/Fd, quenching of chlorophyll fluorescence provided by their binding into chlorophyll–protein complexes (due to pigment–protein interactions); Qm+pi=(Fx-Ft)/Ft, quenching of free pigment fluorescence provided by their incorporation into chlorophyll–protein complexes and in the thylakoids.

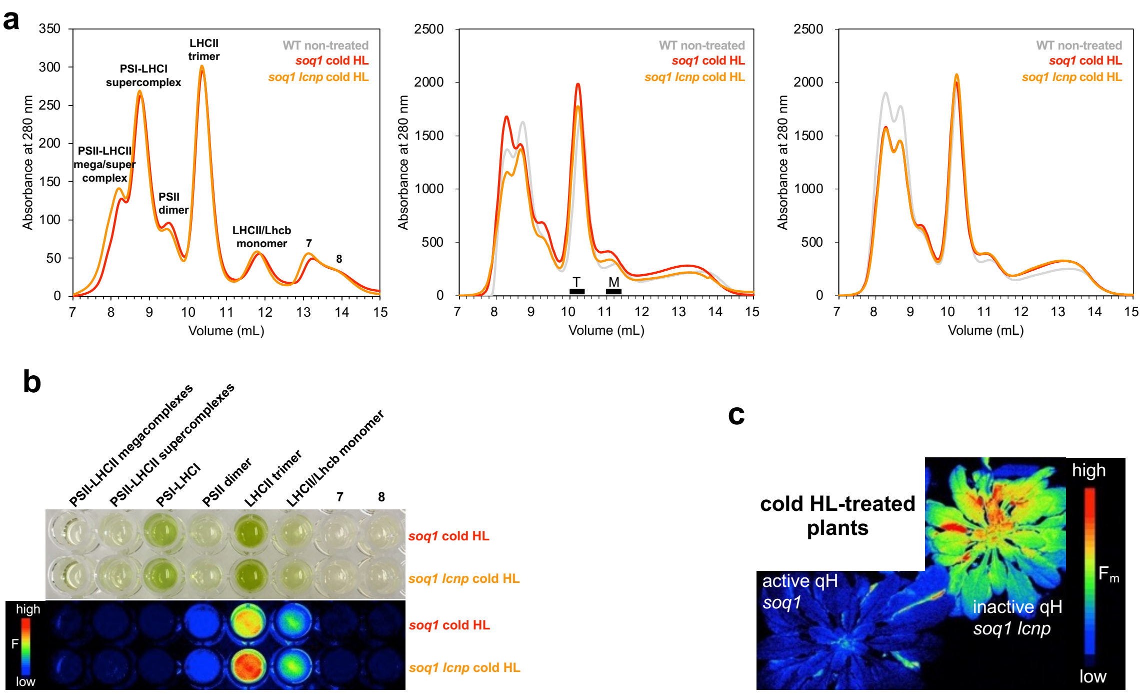


**Fig. S3. qH site is in the LHCII trimer (supports Fig. 2).** Thylakoids were isolated from cold and high light-treated plants and solubilized with 1% $\alpha$-DM at 0.5 mg $mL^{-1}$ chlorophyll concentration for 15 min on ice. Same chlorophyll content (100 µg) was loaded onto the column. **(a)** Gel filtration biological replicate 1 with n=2 plants (left), replicate 2 (middle) and replicate 3 (right) with n = 8 plants from non-treated wild type (WT) (grey), and cold and HL-treated *soq1* (red) and *soq1 lcnp* (orange). Trimer (T) and monomer (M) fractions from replicate 2 used in immunoblot analysis (Fig. S7) are indicated by black bars. Variation in the abundance of the smaller elution volumes (<10 mL) between the three biological replicates is likely due to slight differences in solubilization efficiency (preserving more or less higher orders of PSII-LHCII complexes). **(b)** Top, cropped image of 96-well plate with 200 µl of pooled fraction (three for each) corresponding to peaks annotated in (a). Peaks 7 and 8 contain smaller proteins (composition not analyzed). Bottom, false-colored image of fluorescence yield from these fractions. Fluorescence yield is lower in the LHCII trimer with active qH. Representative from three independent biological experiments is shown; small difference in the LHCII/Lhcb monomer fraction observed here was not significant. **(c)** False-colored image of maximum fluorescence ($F_m$) from representative cold HL-treated plants.

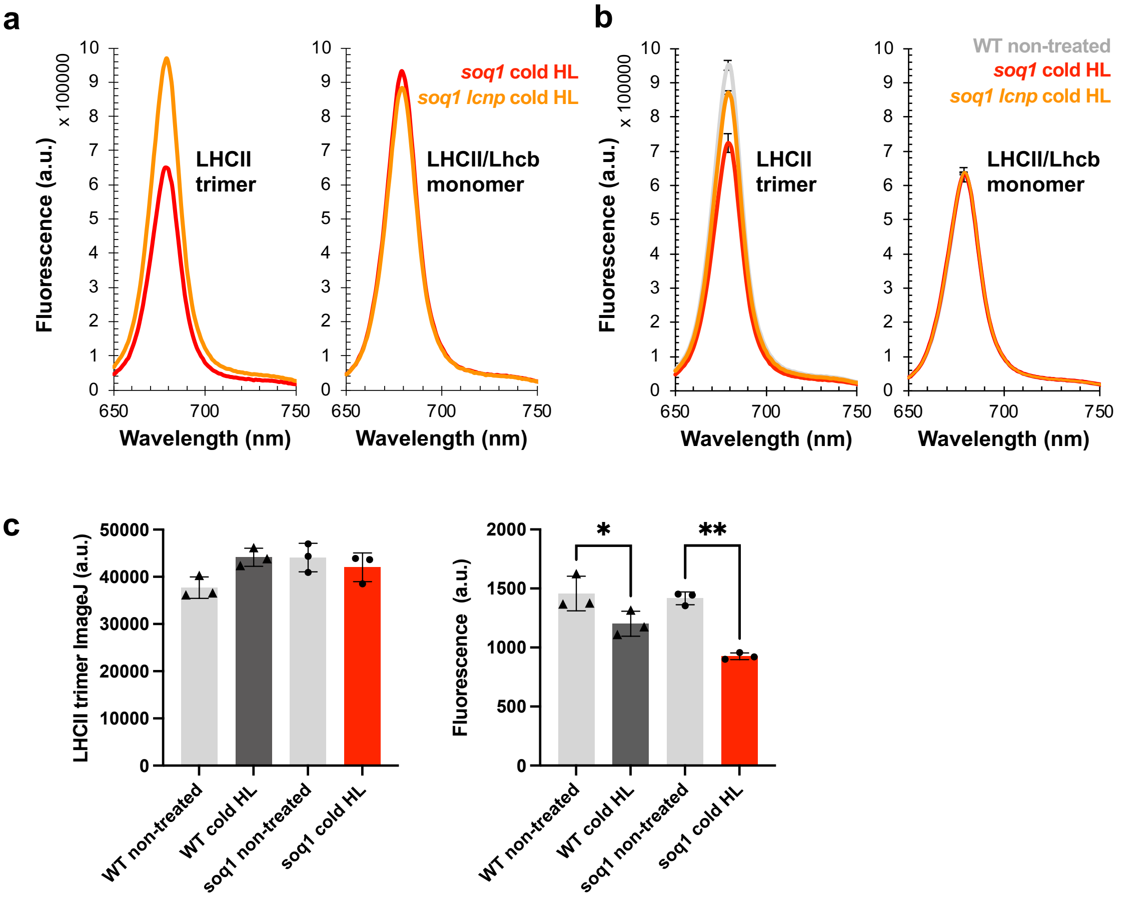


**Figure S4. Fluorescence yield is ~24% lower with active qH in LHCII trimer fraction only (supports Figure 2). (a, b)** Room temperature fluorescence spectra of isolated LHCII trimer and LHCII/Lhcb monomer. Fluorescence emission from 650 nm to 750 nm from samples diluted at same chlorophyll concentration (0.1 µg $mL^{-1}$) with excitation at 625 nm. **(a)** biological replicate 1 (n=2 plants), cold HL-treated 32% difference in yield in trimer with active vs. inactive qH, from upper (U) thylakoid band samples (see Fig. S7a), similar result with lower (L) thylakoid band samples, **(b)** biological replicate 2 (n=8 plants), cold HL-treated 17% difference in yield in trimer. Data represent means ±SD (n=3 technical replicates). **(c)** Quantification of the LHCII trimer pigment-protein band using ImageJ (left) and its chlorophyll fluorescence using SpeedZenII software from JBeamBio (right) from the CN-PAGE shown in Figure 2b. No statistical difference is measured between the samples pigment-protein content. Tukey's multiple comparisons test shows a significant decrease in fluorescence level between WT non-treated and WT cold HL **P*=0.0450 and between *soq1* non-treated and *soq1* cold HL ***P*=0.0011. Data represent means of technical replicates ± SD (n=3 independent loaded gel lanes).

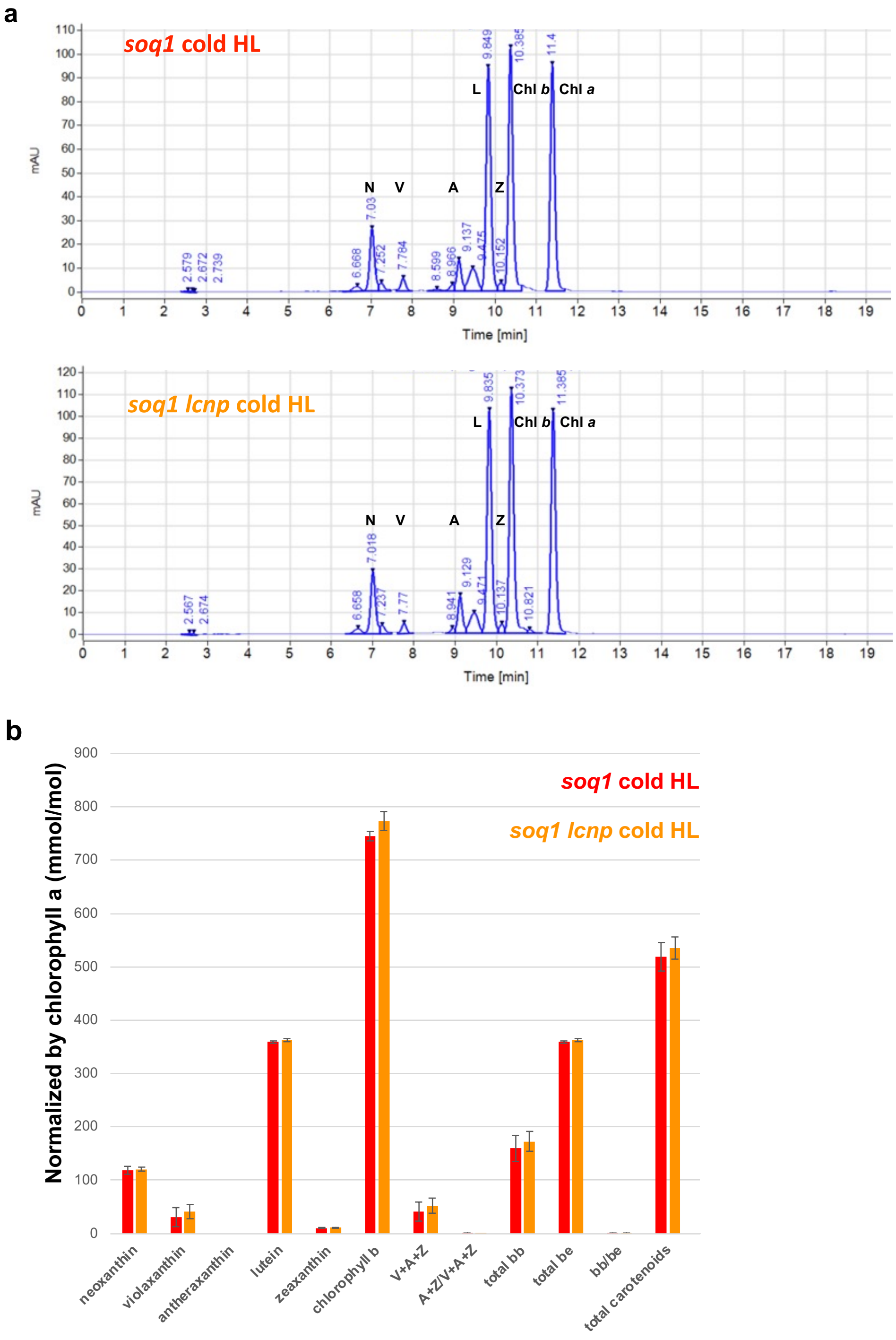


**Fig. S5. No apparent differences in pigment composition and abundance in LHCII trimer fraction. (a)** HPLC traces from LHCII trimer fractions of cold and high light-treated *soq1* and *soq1 lcnp* for 6 h. N, neoxanthin; V, violaxanthin; A, antheraxanthin; Z, zeaxanthin; L, lutein; Chl, chlorophyll. Traces are representative of 2 independent biological replicates each with n = 8 plants. **(b)** Average pigment content normalized to chlorophyll *a*. Data represent means +/- SD (n= 2 independent biological replicates each with n = 8 plants); Compared fractions displayed similar absorbance at 280 nm (Fig. S3a) and had similar Chl concentration (0.25 µg Chl·µ$L^{-1}$) .

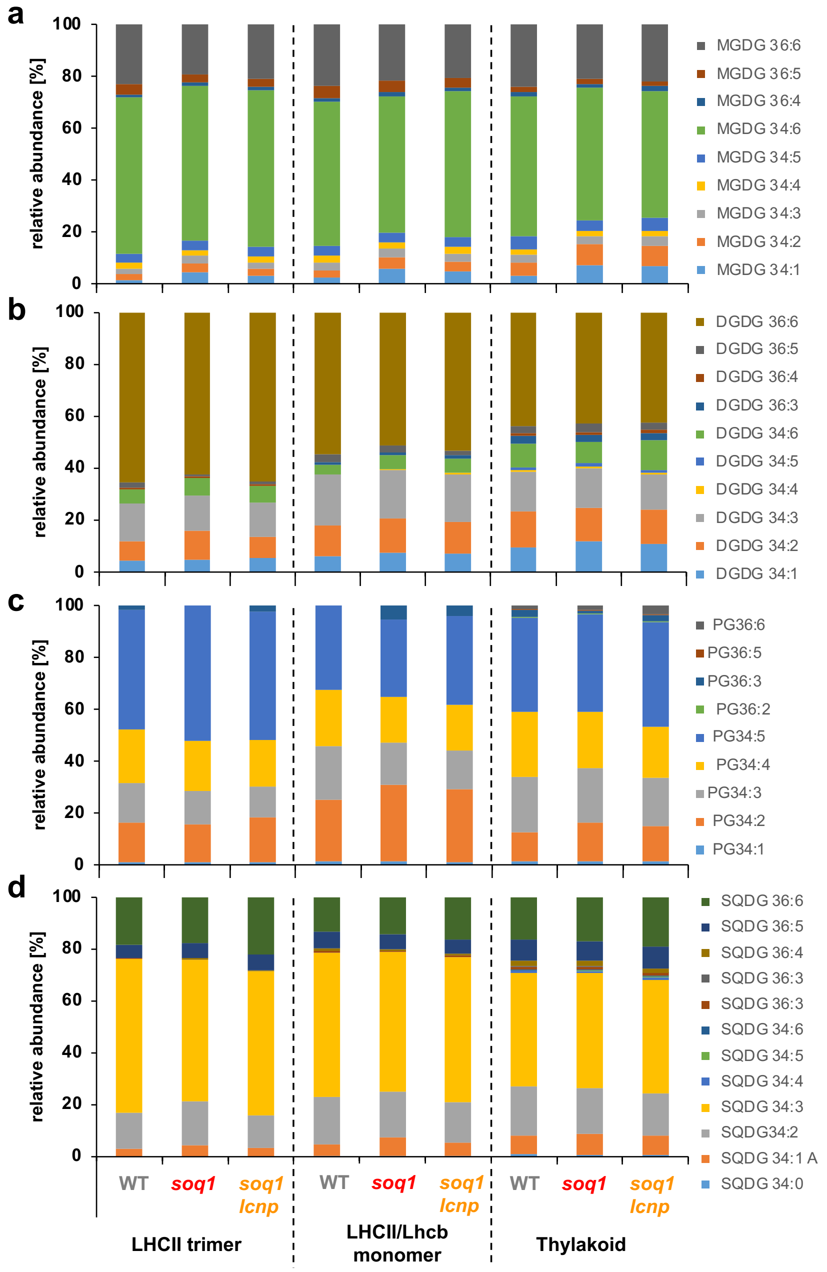


**Fig. S6. No significant differences in lipid composition with active or inactive qH (supports Figure 4).**

Lipid composition of pooled fractions from isolated LHCII trimer from non-treated wild type (WT) or cold HL-treated (replicate 2, n=8 plants) *soq1* (active qH) and *soq1 lcnp* (inactive qH) (see Fig. S3a, middle panel for gel filtration experiment from which fractions were pooled). **(a)** monogalactosyldiacylglycerol (MGDG), **(b)** digalactosyldiacylglycerol (DGDG), **(c)** phosphatidylglycerol (PG) and **(d)** sulfoquinovosyldiacylglycerol (SQDG) species. Relative abundance is calculated per sample based on sum of given species present; absolute quantity normalized to chlorophyll content also did not show significant differences. Small differences in lipid abundance between replicates 2 and 3 could be explained by slight solubilization differences leaving smaller or bigger ring of lipids around the complexes.

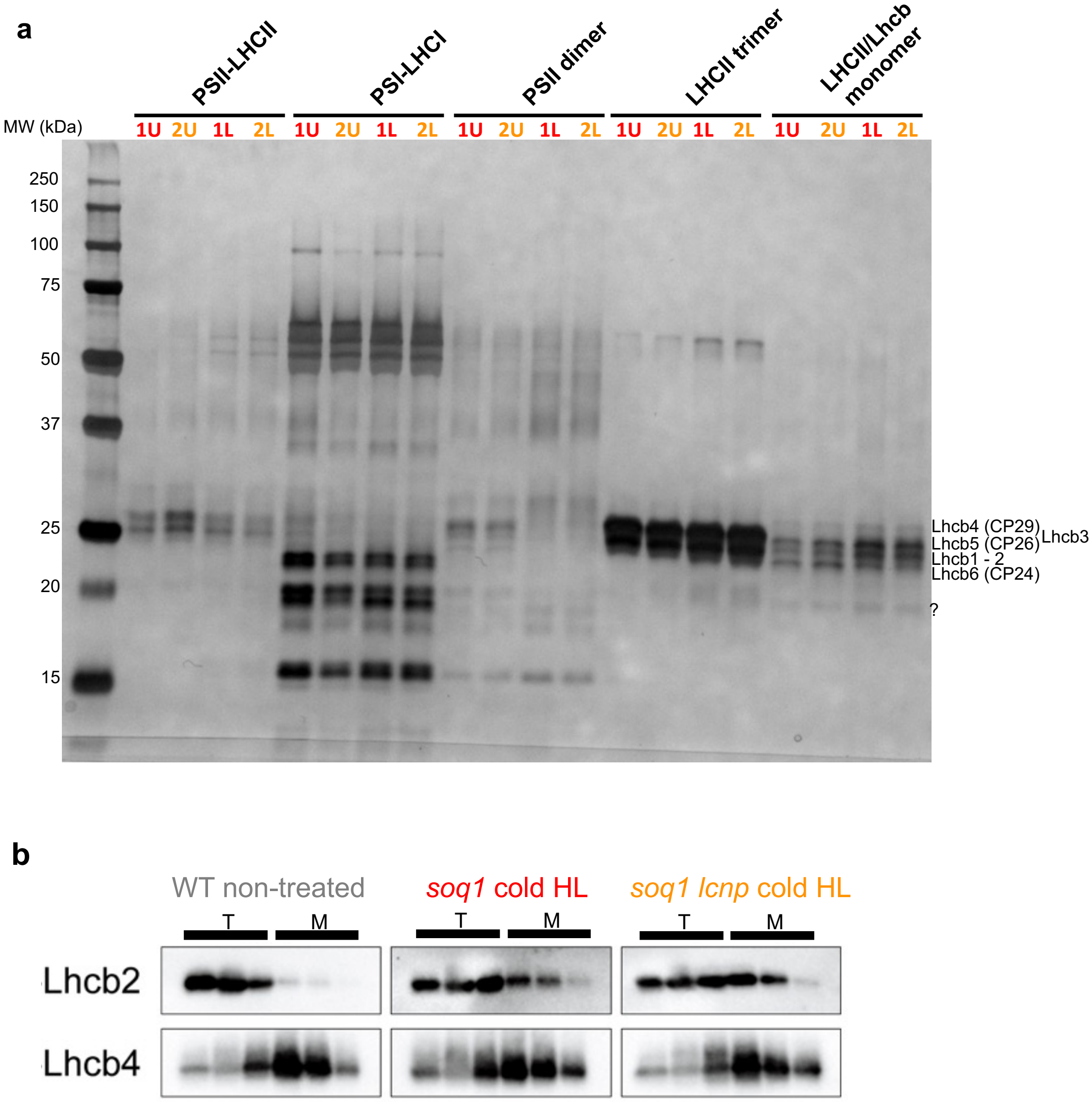


**Fig. S7. Gel filtration fractions contain similar content of proteins with LHCII trimer fraction mainly containing major Lhcbs. (a)** SDS-PAGE followed by silver staining with single or pooled fractions samples from gel filtration experiment displayed in Fig. S3a with active qH, cold HL-treated *soq1* (1) or inactive qH, cold HL-treated *soq1 lcnp* (2). Prior to gel filtration and solubilization, thylakoids were isolated on a sucrose cushion and separated into two fractions from an upper (U) and lower (L) band. The reasoning was that the upper band could be enriched in stroma lamellae (non-appressed membranes) and lower band in grana (appressed membranes), and that this separation could help identify differences between the samples. **(b)** Immunoblot (from replicate 2) from non-treated wild type (WT) (grey), and cold and HL-treated for 6 h *soq1* (red) and *soq1 lcnp* (orange). The sample loaded in the middle lane of the trimer (T) and monomer (M) fractions represents the one used in fluorescence experiments. Samples were loaded at same volume from pooled adjacent fractions (three fractions for each, all nine fractions represented by black bar on Fig. S3a, middle). Antibodies against light-harvesting complex Lhcb2 (major antenna subunit) and Lhcb4 (minor antenna subunit) were used to assess the content of the fractions.

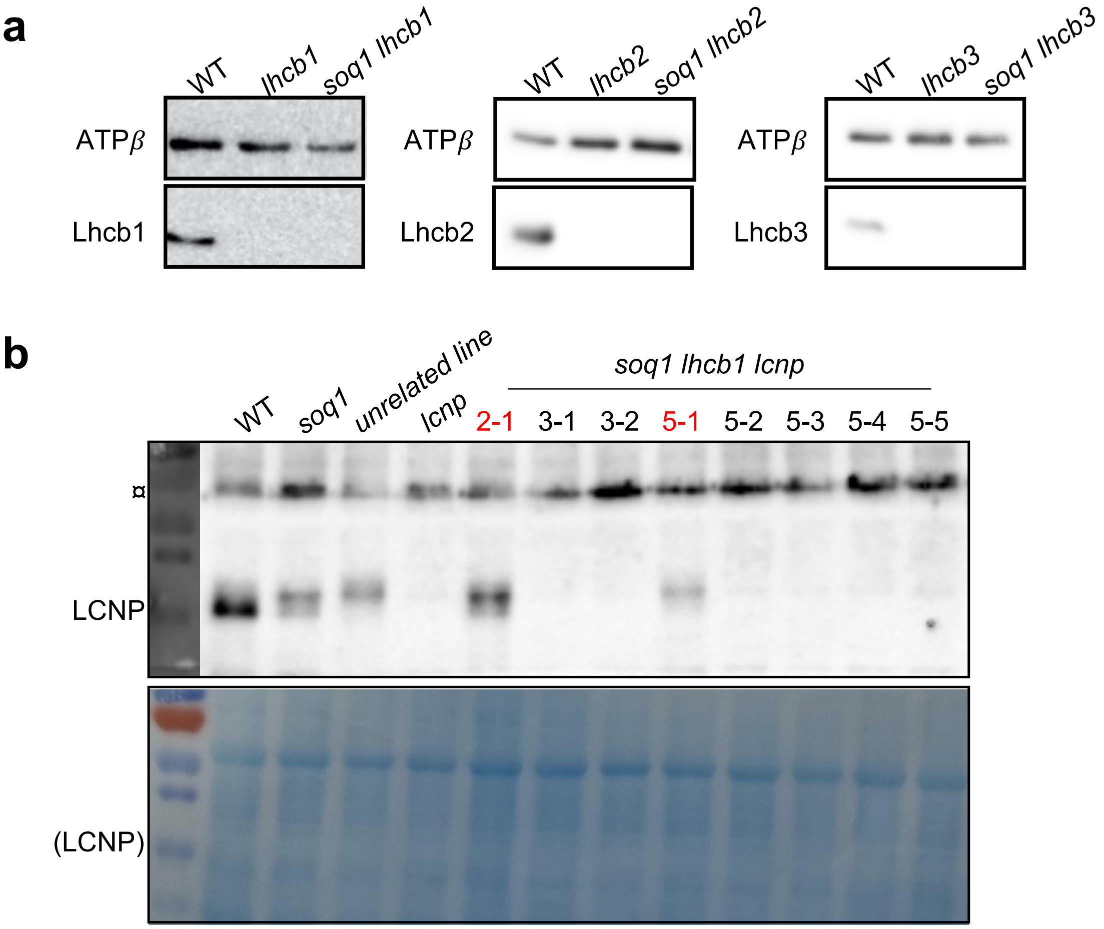


**Fig. S8. qH does not require Lhcb1, or Lhcb2, or Lhcb3 (supports Fig. 5).** Whole cell extract samples from leaves were loaded by equal starting plant material. **(a)** Immunoblot against Lhcb1, Lhcb2 or Lhcb3 in the respective *soq1 lhcb* mutant. ATP*β* is shown as loading control. **(b)** Immunoblot against LCNP in independent individuals from the T1 generation of *soq1 lhcb1 lcnp* lines obtained by CRISPR-Cas9 editing of *LCNP* in a *soq1 lhcb1* background (three individuals were transformed: 2, 3 and 5). The T1 individuals 2-1 and 5-1 marked in red are not knockout (KO) for *LCNP* (a confirmation that the background is *soq1* mutant can be seen by the slower mobility of LCNP) whereas the other individuals are KO for *LCNP*. Individuals 3-1, 3-2 and 5-5 were used for NPQ, $F_o$ and $F_m$ measurements shown in Fig. 5. Symbol (¤) indicates nonspecific band detected by the anti-LCNP antibody. Coomassie blue is shown as loading control and (LCNP) indicates the size where LCNP migrates.

**Table S2. Summary of fluorescence lifetime from leaves, isolated thylakoids, LHCII trimer, LHCII/Lhcb monomer and PSII dimers (supports Fig. 1, 3).** Average fluorescence lifetimes (ns) determined by time-correlated single photon counting from cold and high light-treated plants for 6 h at 6°C and 1500 µmol photons $m^{-2}$ $s^{-1}$ | and from non-treated plants.

<table>
<tr><th>Sample type</th><th>soq1 roqh1</th><th>soq1</th><th>WT</th><th>soq1 (roqh1) lcnp</th></tr>
<tr><td>Leaves</td><td>| 0.10 ± 0.023</td><td>0.13 ± 0.01 | 1.50 ± 0.02</td><td>0.34 ± 0.10 | 1.50 ± 0.05</td><td>0.47 ± 0.11 | 1.42 ± 0.12</td></tr>
<tr><td>Thylakoids</td><td>| 0.18 ± 0.002</td><td>0.30 ± 0.06 | 1.21 ± 0.11</td><td>0.66 ± 0.04 | 1.14 ± 0.09</td><td>0.60 ± 0.03 | 1.10 ± 0.08</td></tr>
<tr><td>LHCII trimers</td><td></td><td>2.64 ± 0.11 |</td><td>| 3.09 ± 0.06</td><td>3.32 ± 0.17 |</td></tr>
<tr><td>LHCII/Lhcb monomers</td><td></td><td>2.58 ± 0.11 |</td><td>| 2.87 ± 0.08</td><td>2.61 ± 0.07 |</td></tr>
<tr><td>PSII dimers</td><td></td><td>1.14 ± 0.09 |</td><td>| 1.22 ± 0.09</td><td>1.19 ± 0.12 |</td></tr>
</table>

**Table S3. Averages of amplitudes and time components for the fits of time-correlated single photon counting data on leaves, thylakoids and isolated complexes.** Each emission decay at 680 nm upon excitation at 420 nm was fit to a tri-exponential model without constraining any specific kinetic component, and an amplitude-weighted average fluorescence lifetime ($\tau_{avg}$) was calculated.

| Sample type | Line | A1 (%) | τ1 (ns) | A2 (%) | τ2 (ns) | A3 (%) | τ3 (ns) | τavg (ns) |
|---|---|---|---|---|---|---|---|---|
| Leaves (non-treated) | WT | 15 | 0.1 | 27 | 0.9 | 58 | 2.1 | 1.5 |
| | *soq1* | 13 | 0.1 | 26 | 0.8 | 61 | 2.1 | 1.5 |
| | *soq1 roqh1 lcnp* | 13 | 0.2 | 31 | 0.9 | 56 | 2.0 | 1.4 |
| | *soq1 roqh1* | **67** | 0.1 | 32 | 0.2 | **1** | 0.8 | **0.1** |
| Leaves (cold HL-treated) | WT | 52 | 0.1 | 37 | 0.5 | 10 | 1.1 | 0.3 |
| | *soq1* | **66** | 0.1 | 33 | 0.2 | **1** | 0.8 | **0.1** |
| | *soq1 roqh1 lcnp* | 35 | 0.1 | 53 | 0.5 | 12 | 1.2 | 0.5 |
| Thylakoids (non-treated) | WT | 29 | 0.1 | 27 | 0.8 | 44 | 2.0 | 1.1 |
| | *soq1* | 26 | 0.1 | 25 | 0.8 | 49 | 2.0 | 1.2 |
| | *soq1 roqh1 lcnp* | 27 | 0.1 | 26 | 0.7 | 47 | 1.8 | 1.1 |
| | *soq1 roqh1* | **68** | 0.1 | 31 | 0.3 | **1** | 2.5 | **0.2** |
| Thylakoids (cold HL-treated) | WT | 38 | 0.1 | 42 | 0.7 | 21 | 1.6 | 0.7 |
| | *soq1* | **57** | 0.1 | 36 | 0.4 | **6** | 1.5 | **0.3** |
| | *soq1 roqh1 lcnp* | 33 | 0.1 | 51 | 0.7 | 15 | 1.5 | 0.6 |
| LHCII trimers (cold HL-treated) | *soq1* | **18** | 0.1 | 9 | 0.5 | **73** | 3.6 | **2.7** |
| | *soq1 lcnp* | 7 | 0.1 | 8 | 0.8 | 85 | 3.8 | 3.3 |
| LHCII/Lhcb monomers (cold HL-treated) | *soq1* | 18 | 0.1 | 12 | 0.9 | 70 | 3.5 | 2.6 |
| | *soq1 lcnp* | 16 | 0.1 | 13 | 0.8 | 71 | 3.5 | 2.6 |
| PSII dimers (cold HL-treated) | *soq1* | 53 | 0.1 | 19 | 0.6 | 28 | 3.5 | 1.1 |
| | *soq1 lcnp* | 51 | 0.1 | 19 | 0.6 | 30 | 3.4 | 1.2 |

**Table S4.** Primer list used in this study.

| | |
|---|---|
| *LHCB1.1* Forward | AGGCAGTTTGGTTCAAGGCT |
| *LHCB1.1* Reverse | GCCTCTACAACGGAGTGAACC |
| *LHCB1.2* Forward | TCAGCTGATCCCGAGACAT |
| *LHCB1.2* Reverse | CTGAAAGTCTCAAACCATCAC |
| *LHCB1.3* Forward | GTGCTGCACTACTCAACC |
| *LHCB1.3* Reverse | TCACACTCACGAAGCAAAGACTG |
| *LHCB1.4* Forward | TAGGCTGCGTTTTCCCTGAG |
| *LHCB1.4* Reverse | TCCAAGATGCAACAAACCGGA |
| *LHCB1.5* Forward | AACCATGGCTTTGTCCTCCC |
| *LHCB1.5* Reverse | CTCTGCTCTCATTCACATAATAAGCAG |
| *LHCB2.1* Forward (SALK_005774C) | AAGTTTCAACTGAGCCAAAAAG |
| *LHCB2.1* Reverse (SALK_005774C) | TTGGTACCAGATGCTTTGAGG |
| *LHCB2.2* Forward | GCATTGTAATGGCTCATGACC |
| *LHCB2.2* Reverse | CGTAGTCTCCGGGGTATTCTC |
| *LHCB2.3* Forward | TCCAACACTCTTCTTTCGCTG |
| *LHCB2.3* Reverse | ACGTACTGCCAAGTTGCTAGC |
| *LHCB3* Forward (SALK_036200C) | TTAAATGGGCAACCAGAAAAG |
| *LHCB3* Reverse (SALK_036200C) | ACCCAACGGGTCAAAGTATTG |
| LBa1 (SALK line genotyping) | TGGTTCACGTAGTGGGCCATCG |
| *SOQ1* Forward (dCAPS) (Malnoë et al. 2018) | GAAGTGGTTTCTTTTGTACAATTCTGCA |
| *SOQ1* Reverse (dCAPS) (Malnoë et al. 2018) | CAATACGAATAGCGCACACG |
